# Evaluating the predictive accuracy of ion channel models using data from multiple experimental designs

**DOI:** 10.1101/2024.08.16.608289

**Authors:** Joseph G. Shuttleworth, Chon Lok Lei, Monique J. Windley, Adam P. Hill, Simon P. Preston, Gary R. Mirams

**Affiliations:** Centre for Mathematical Medicine & Biology, School of Mathematical Sciences, University of Nottingham, UK; Institute of Translational Medicine, Faculty of Health Sciences, University of Macau, Macau, China; Department of Biomedical Sciences, Faculty of Health Sciences, University of Macau, Macau, China; Computational Cardiology Laboratory, Victor Chang Cardiac Research Institute, Darlinghurst, New South Wales, Australia; School of Clinical Medicine, Facility of Medicine and Health, University of New South Wales Sydney, New South Wales, Australia

## Abstract

Mathematical models are increasingly being relied upon to provide quantitatively accurate predictions of cardiac electrophysiology. Many such models concern the behaviour of particular subcellular components (namely, ion channels) which, together, allow the propagation of electrical signals through heart-muscle tissue—namely, the firing of action potentials. In particular, I_Kr_, a voltage-sensitive potassium ion-channel current, is of interest owing to its central pore’s propensity for blockage by various small molecules. We use newly collected data obtained from an ensemble of voltage-clamp experiments to validate the predictive accuracy of various dynamical models of I_Kr_. To do this, we fit models to each protocol individually, and quantify the error in the resultant model predictions. This allows the comparison of predictive accuracy for I_Kr_ models under a diverse collection of previously unexplored dynamics. Our results highlight heterogeneity between parameter estimates obtained from different cells, suggesting the presence of latent effects not yet accounted for in our models. This heterogeneity has a significant impact on our parameter estimates and suggests routes for model improvement.

## 1 Introduction

### 1.1 I_Kr_ and its role in cardiac electrophysiology

Present throughout the human body, ion channels are protein structures embedded in the cell membrane which play an important role in the transmission and reception of electrical signals. This is especially true of the production of cellular action potentials in heart-muscle cells (cardiomyocytes) [1]. Ion channels mediate the flow of specific species of ions into and out of the cell. The focus of this paper, K_V_11.1, is a potassium ion channel which opens and closes in response to voltage signals (that is, a voltage-sensitive potassium ion channel). In heart-muscle cells (cardiomyocytes) there are a large number of K_V_11.1 channels, the combined current through which is known as the *rapid delayed rectifier potassium current* and denoted by I_Kr_ [2]. The block of this current by small molecules is associated with dangerous changes to the heart’s rhythm (arrhythmia) [3]. Consequently, K_V_11.1 is a key focus of drug safety assays [4]. For this purpose, accurate mathematical models of the baseline behaviour of I_Kr_ (among other ion-channel currents) are desirable, allowing the impact of drug-channel interactions to be quantified and used to classify proarrhythmic risk [5, 6]. Whilst our work here is focused on hERG1a cell lines rather than cardiomyocytes, we use I_Kr_ as a shorthand for the recorded currents. The methods herein are presented such that they may be applied broadly to other macroscopic, voltage-gated ion channel currents.

Typically, models of macroscopic ion-channel currents are built, trained, and validated using data collected from *patch-clamp electrophysiology* experiments [7, 8]. We adopt this approach in this paper, performing whole-cell voltage-clamp experiments on a high-throughput, automated patchclamp platform. These experiments allow I_Kr_ to be measured whilst the transmembrane potential, *V*_m_, is manipulated. Such experiments can produce a wealth of information-rich data [9], which may be used to fit mathematical models of macroscopic ion channel currents [10, 11, 12]. A diagram of such an experiment is shown together with an equivalent electrical circuit in Figure 1. We use the resulting data to train and validate I_Kr_ models, as described below.

**Figure 1:**
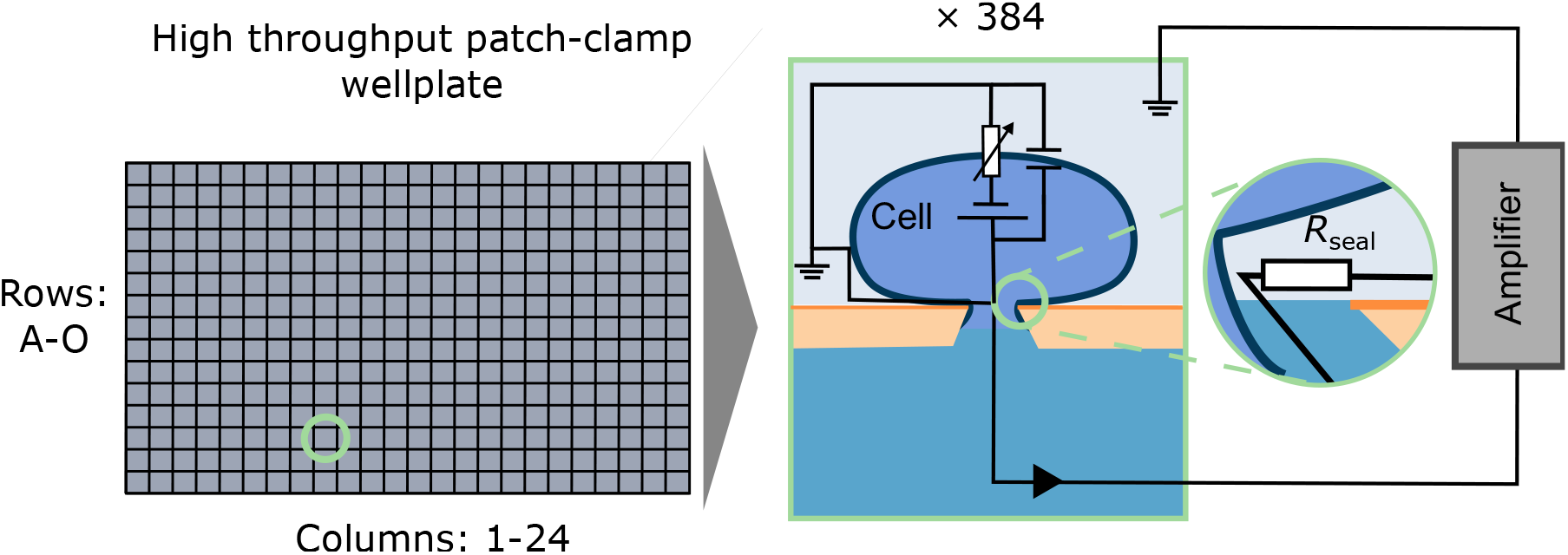
A diagram of a patch-clamp experiment performed on a high-throughput, automated patch-clamp platform. A seal is formed between the plate and the cell. Automatically applied pressure is then used to puncture the cell membrane such that an electrical current flows from the inside of the cell, through the membrane, to the amplifier where it is recorded. Figure modified from [13].

Whilst mathematical models are increasingly being used to quantify the impact that drugs have on I_Kr_ and other ion-channel currents [6], many conflicting, yet plausible, mathematical models are suggested in the literature [14, 15]. Typically, these models are *Markov models*[10], including different numbers of parameters and differences in the number of states and how they are connected (the *model structure*). Regarding the choice of model structure, Mangold *et al*. enumerated many thousands of possible model structures for Markov models of the fast sodium current, *I*_Na_ and the fast-transient outward potassium current [16]. Many of these structures may also yield plausible models of I_Kr_ [9], albeit with drastically different parameter values. Note that other models, such as so-called Hodgkin-Huxley models, may be expressed as Markov models, even if they were not originally presented this way [10, 17, 18]. However, it is not clear which model (or models) provide the most accurate description of I_Kr_ [9], or which are the most suitable for use in drug-binding assays or inclusion in whole-cell action-potential models [18, 19]. By modelling the dynamics of these models under various protocols, and performing extensive validation of our models’ predictive accuracy, we aim to select the most suitable model structures, and, as a result, obtain more accurate predictive models of I_Kr_.

Though probabilistic models may be used to study ion channels [20] (especially for models of single channels [21]), Markov models of I_Kr_ are typically implemented deterministically as systems of ordinary differential equations (ODEs) [10]. In all the models we consider, only a single ‘open’ conformation allows current to flow through the channel, and so the I_Kr_ current is proportional to the fraction of channels in this conformation. Numerous such models of I_Kr_ are suggested in the literature, with various contradicting model structures; they disagree on the number of conformational states, which transitions between states are possible, and certain symmetries between the rates at which transitions occur [14]. We aim, therefore, to develop the methodology necessary to select the most accurate from a pool of candidate models.

Using synthetically generated data, we have previously shown that we are able to identify a correct model structure from amongst a pool of candidates, resulting in improved predictive accuracy [22]. Here, we adapt and apply this methodology to newly collected experimental data using a wider range of experimental designs. In particular, we apply a diverse range of voltage-clamp *protocols* (that is, the user-defined voltage signals that comprise our experimental design) to a selection of cells and record the resulting currents simultaneously. However, the extension of this work to real data introduces additional complications—not least of which is the fact our approximate mathematical models are incapable of fully recapitulating the underling *data-generating process* (DGP) [23]. Nevertheless, this newly collected data allows us to compare the predictive accuracy of a collection of literature I_Kr_ models.

The variability in parameter estimates obtained by fitting models to data from different cells has been explored previously in the literature [24]. It is unclear to what extent this is the result of underlying biological variability or other non-biological factors affecting the recorded current (experimental artefacts). We consider the variability of our parameter estimates (each taken from a given well and protocol) in Section 6. Where we show that our parameters estimates depend, not only on the voltage protocol used, but on the particular cell/well the data was obtained from. This provides insight into the nature of the discrepancy between our mathematical models and the DGP. This analysis suggests the presence of latent well-dependent effects, not yet accounted for in our mathematical models.

## 2 Mathematical models of I_Kr_

Each of the four Markov models we consider is an ODE-based model with a *governing equation*,

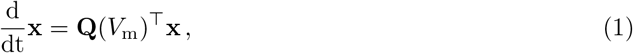

where **x** is a *state-variables* vector which describes the portion of channels in each of the model’s conformational states, **Q** is a voltage-dependent transition rate matrix [25] such that the element *Q*_*i*,*j*_ is the transition rate between the model’s *i*^th^ and *j*^th^ states, and *V*_m_ is the *transmembrane potential* that is, the potential difference between the inside and outside of the cell membrane. These states are mapped to our observables, via an *observation function* of the form,

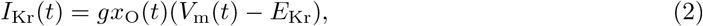

where *E*_Kr_ is the *reversal potential, g* is the maximal conductance, and *x*_O_ is the state in the vector **x** representing the open channel conformation.

Typically, the model’s reversal potential, *E*_Kr_, is set to the Nernst potential which may be calculated as,

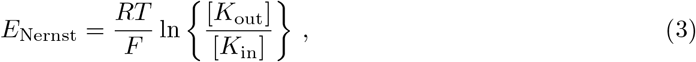

where [*K*_out_] denotes the extracellular potassium concentration, [*K*_in_] denotes the intracellular potassium concentration, *R* is the gas constant, *F* is Faraday’s constant, and *T* = 298.15 K = 25^*o*^C is the temperature at which our experiments were performed [26]. This is the transmembrane potential at which, according to the model, there is no force driving K^+^ ions through the channel (see Equation 2). Using the known potassium concentrations of our intracellular and extracellular solutions, we find *E*_Nernst_ ≈ −90 mV.

We assume that the initial state vector, **x**(0), lies at the governing equation’s unique global equilibrium point. Such an equilibrium point is guaranteed to exist for our choice of models [27, 10]. The assumption that the model is at equilibrium when *t* = 0 is made because the cell is left to equilibrate before each protocol (that is, before each *sweep* is recorded). During this time, the command voltage, *V*_cmd_ is held at the holding potential, −80 mV, and we can compute the resulting steady state which depends on the model parameters, ***θ*** [10].

Various Markov models, each characterised by a different choice of **Q** can be used here. Note that the number of states in the model, *N*, may also vary, meaning that the length of the state-variable vector, **x** ∈ ℝ^*N*^, may differ between models. For each model, transition rates, *Q*_*i*,*j*_ are dictated by our model parameters. Typically, we have rates of the form *Q*_*i*,*j*_ = *A* exp{±*bV* } where *A* and *b* are model parameters (such as *p*_1_ and *p*_2_ in the Beattie model [9]). The Wang model [28], however, contains two voltage-independent transition rates, *k*_*f*_ and *k*_*b*_, which are, themselves, scalar model parameters. Also, for each model, the maximal conductance, *g*, is fitted as an additional model parameter, acting as a scaling factor (see Equation 2).

We consider the four models shown in Figure 2: the Beattie model [9]; the Kemp model [29]; the Wang model [28]; and a simple, three-state model which we refer to as the closed-open-inactive (C-O-I) model. These models differ in the number of states (that is, *N*) and parameters (those which determine transition rates), and in the existence of a path between the inactivated state (**I**) and the closed states (**C/C1/C2/C3**) that avoids the open state, **O**. However, each model shares the same form, satisfying Equations 1 and 2. Whilst the reversal potential is treated as a known constant, the maximal conductance, *g*, like our transition-rate parameters, is fitted independently for each individual sweep.

**Figure 2:**
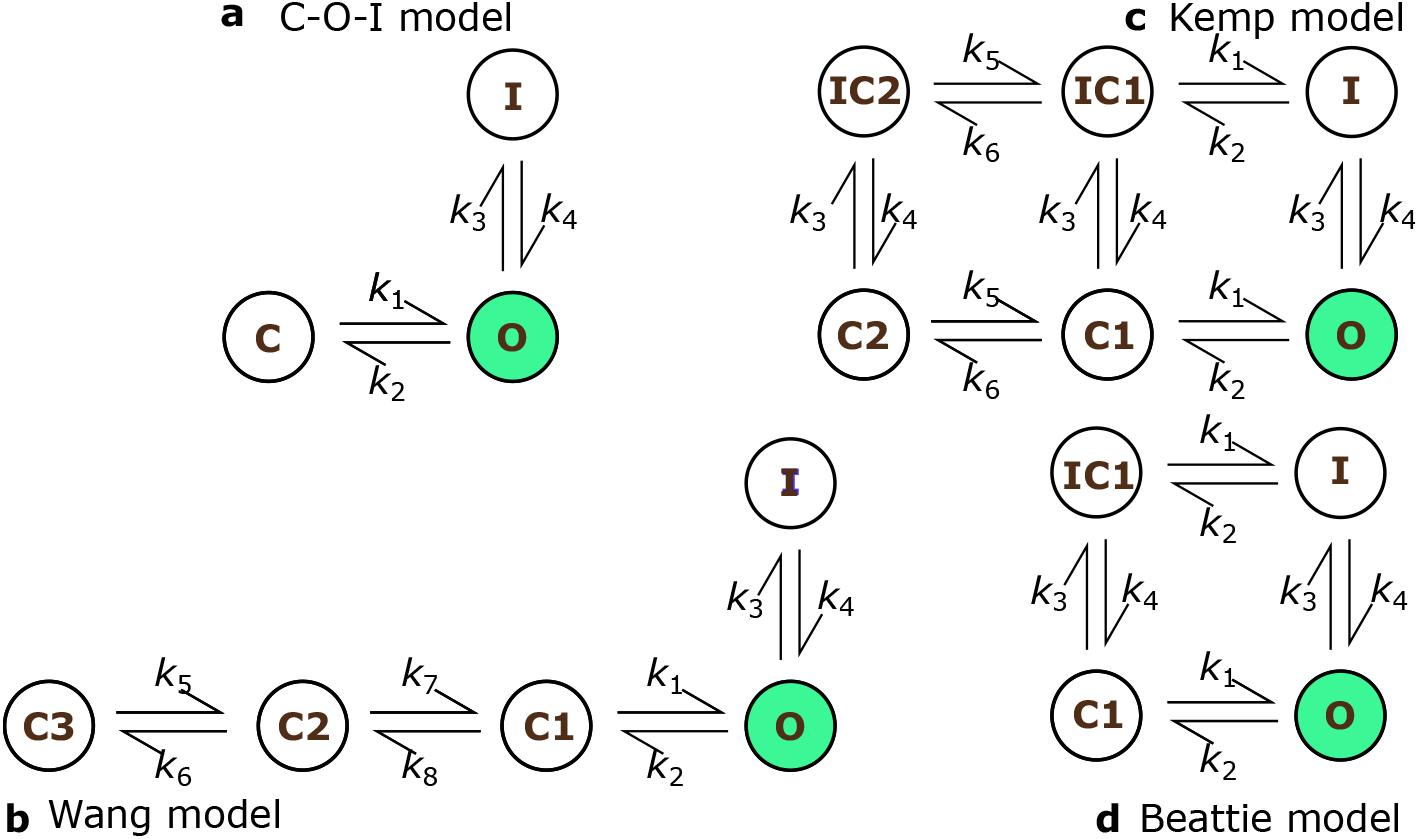
The four model structures used. Note that the **C1** to **C2** transition in the Wang model is constant, that is, determined by a single model parameter, whereas all other transition rates are parameterised by exactly two parameters. All rates with even indices have rates of the form, *k*_2*i*_ = *a* exp{*bV*_m_} and all odd numbered rates are of the form, *k*_2*i*+1_ = *a* exp{-*bV*_m_}, with the exception of the Wang model’s *k*_5_ and *k*_6_ which are both constant rates[28].

To fit a given model, we assume that the data was generated using the given model structure with some unknown parameter set, ***θ***. Then, to fit our model, we that our observations, *z*_*i*_, are subject to additive, independent and identically distributed (IID) Gaussian errors, *ε*_*i*_ such that,

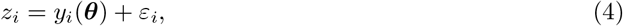

where *y*_*i*_(***θ***) is the observable corresponding to the *i*^th^ observation, which depends on the model parameters, ***θ***. Then, we compute the maximum likelihood estimate (MLE), 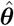, by finding the parameter set which minimises 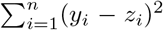, where *n* is the number of observations. For this model, maximum likelihood estimation is equivalent to nonlinear least-squares regression. After computing a parameter estimate for each sweep and for each model, we use these parameter estimates to compute predictions for the remaining protocols.

We assume that our data arises from an *ideal* patch-clamp setup. That is, we assume that at any time, *t*, the membrane voltage is exactly the command voltage, albeit with a possibly non-zero systematic voltage error that is,

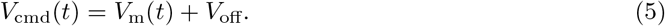

This voltage-offset is included to explain the discrepancy between *E*_Nernst_ and *E*_obs_ as discussed in the Electronic Supplement (Appendix D). Secondly, we assume that the current we observe during the experiment is exactly a combination of I_Kr_ and our linear-leak current,

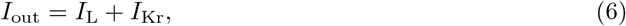

where *I*_L_ is the leak current satisfying,

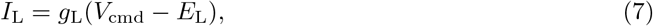

where *g*_L_ is the leak conductance and *E*_L_ is the reversal potential of the leak current. These two parameters are fitted during *postprocessing* (before our Markov-model parameters are fitted) as described in Section 5.

## 3 Experimental methods

### Cell culture and harvesting

hERG channels stably expressed in CHO cells were purchased from the American Type Culture Collection (ATTC reference PTA-6812). CHO cells were maintained in Hams Nutrient mix media (ThermoFisher Scientific, Waltham, USA) supplemented with 5% fetal bovine serum (Merck Life Science, Melbourne, VIC, Australia). Cells were housed in a 37°C, humidified incubator at 5% CO_2_. Cells were passaged every 2–3 days and harvested for experiments 48–72 hours after passaging. Prior to harvesting, cells were grown in t150 or t175 tissue culture flasks to a confluency of 60-80%. Confluent cells were washed twice with PBS (Mg/Ca^2+^ free, ThermoFisher Scientific, Waltham, USA) and incubated with Accumax (Merck Life Science) at 37°C for 4–5 minutes to enable cell detachment. Cells were incubated for an additional 5 minutes at 4°C following the addition of cold SyncroPatch recording solution (see below) to allow for membrane recovery prior to manual agitation and removal of cells from the flask. Cells were centrifuged for 5 minutes at 250 g, the supernatant removed and resuspended in divalent free SyncroPatch solution (see below) to a density of 250–500 thousand cells/ml. The cellular suspension was incubated for an additional 30–60 minutes at 4 °C and finally transferred to the shaking SyncroPatch cell platform, which was maintained at 10°C throughout the experiment.

### High throughput patch clamp setup and solutions

To get the highest quality recordings it is important to note that we used fluoride-free plates and solutions [30], as in the first attempts in fluoride-containing pilot experiments we saw larger non-linear time dependent leak currents that were difficult to isolate and remove [13].

Patch clamp experiments were performed on the SyncroPatch 384PE (Nanion, Munich, Germany). Nanion, 1 hole, medR FF (fluoride free, 4–4.5 MΩ) and NPC-384T L-type (standard, 3– 5 MΩ) SyncroPatch plates were used to run 384 whole cell, patch clamp recordings in parallel. Cell catching, sealing, whole cell breakthrough and capacitance compensation procedures were automated by the SyncroPatch. According to Nanion’s fluoride-free chip procedures, fluoride-free experimental plates were pre-treated with 0.5 mM NaOH prior and washed three times with water and divalent free solution as part of the automated program. In addition, to improve the success rate with respect to series resistance the membrane perforator, Escin (15 μM, Merck Life Science) was added to the internal solution and washed out with Escin free internal solution following the whole cell pressure pulse step. Experiments were performed at ambient temperature (25±1°C). The standard, fluoride containing SyncroPatch internal solution contained, 110 mM KF, 10 mM KCl, 10 mM NaCl, 10 mM HEPES and 10 mM EGTA. The fluoride-free internal solution contained, 120 K gluconate, 10 mM KCl, 10 nM NaCl, 10 mM HEPES and 5 mM EGTA. Both internal solutions were adjusted to pH 7.2 with KOH. The divalent free solution used for the cell suspension and initial filling of the SyncroPatch plates contained, 140 mM NaCl, 4 mM KCl, 5 mM glucose and 10 mM HEPES for all experiments. Seal enhancing solutions employed to improve cell-to-plate seal performance contained, 80 mM NaCl, 60 nM NMDG-Cl, 10 mM CaCl_2_, 1mM MgCl_2_, 5 mM glucose and 10 mM HEPES for standard, fluoride containing experiments, while fluoride free seal enhancer contained, 140 mM NMDG-Cl, 4 KCl, 4 mM CaCl_2_, 1 mM MgCl_2_, 5 mM glucose and 10 mM HEPES. The recording solution for fluoride and fluoride free experiments contained, 80 mM NaCl, 60 mM NMDG-Cl, 4 mM KCl, 2 mM CaCl_2_, 1 mM MgCl_2_, 5 mM glucose and 10 mM HEPES. All solutions, except for internal, were adjusted to pH 7.4 with NaOH. All chemicals, unless otherwise stated, were purchased from Merck Life Science. Individual liquid junction potentials where calculated for experiments with and without fluoride and adjusted in SyncroPatch protocols accordingly.

### Pharmacological Isolation of I_Kr_ current

After applying each voltage protocol to our cells, we add *dofetilide* at 1 *μ*M (a concentration known to almost fully block I_Kr_) and repeat each protocol in the same order as shown in Figure 4. By performing leak correction and subtracting the post-drug leak-corrected trace from the pre-drug leak-corrected trace, we are able to isolate I_Kr_ with minimal contamination from any endogenous or other currents. These postprocessing methods are explained in Section 5.

**Figure 3:**
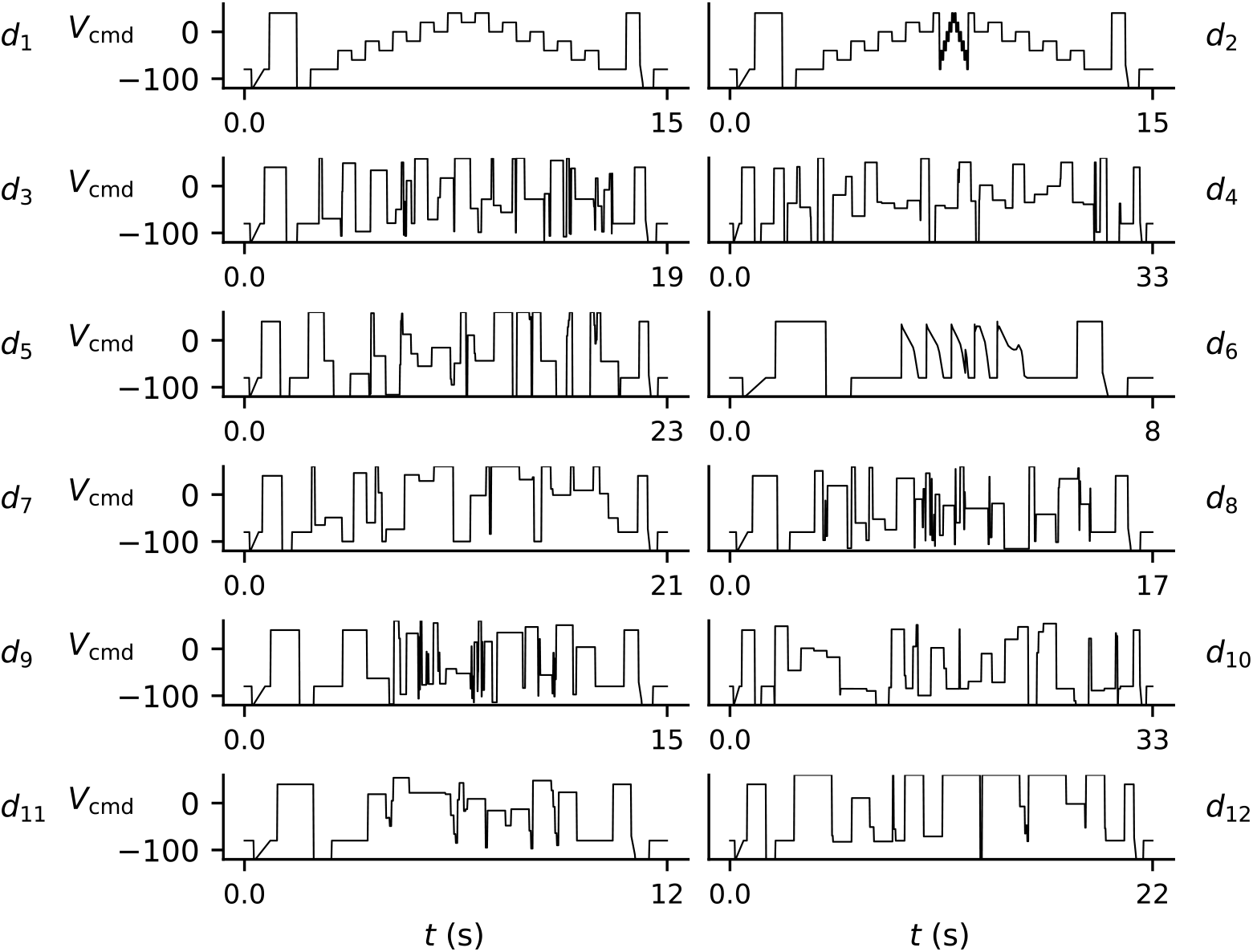
The protocols used in our experiment, shown in the order that they are applied. Common features present at the start and end of each protocol are used for postprocessing. All protocols are repeated exactly once except *d*_1_, Lei *et al*.’s *staircase* protocol. All protocols are used for model validation except *d*_6_ which is used only for validation.

**Figure 4:**
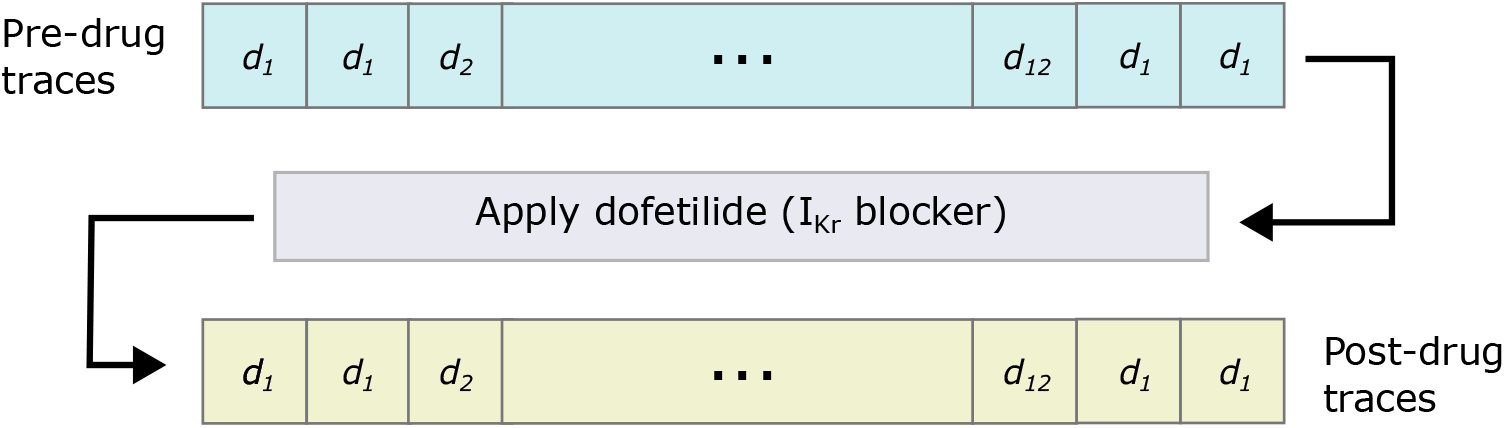
The order in which the protocols were performed. First two sweeps of *d*_1_ (the *staircase* protocol) are performed, then a single sweep of each of the other protocols are performed before two final sweeps of *d*_1_. Then, after the addition of 1 *μ*M dofetilide (which should provide a specific I_Kr_ block), this sequence of protocols is repeated once more. This allows the subtraction of post-drug traces from pre-drug traces, which mitigates the presence of endogenous (non-I_Kr_) currents and any other artefacts.

Dofetilide (Merck Life Science) was prepared as 10 mM stocks in 100% DMSO. Drug stocks were used immediately or stored at −20°C in glass opaque vials, in small aliquots for single use only (Merck Life Science). In the latter case, stocks were thawed immediately prior to experiments, vortexed and prepared in recording solution to the appropriate concentrations in glass vials. Drug solutions were then transferred to Teflon SyncroPatch plates for automated addition to the SyncroPatch plate following the recording of hERG channel currents in drug free recording solutions.

## 4 Design of voltage-clamp protocols

A range of *information-rich* voltage protocols were applied sequentially to each well. The differences lie in the specified ‘command voltage’, *V*_cmd_, that is, the voltage the amplifier is instructed to clamp the membrane potential to at each time point during the experiment. These protocols were developed using a range of techniques, as detailed in Lei *et al*. [31]. We will briefly describe the rationales for their designs again here.

Of particular importance is the *staircase* protocol, *d*_1_, of which we perform four repeats. These repetitions allow us to ensure that the cell’s response to *V*_cmd_(*t*) remains constant over the course of the experiment (see Appendix B of the Electronic Supplement for more detail). The remaining protocols are performed once each.

Some protocols were designed via numerical optimisation with respect to various objective functions. In particular: protocols *d*_3_, *d*_8_ and *d*_9_ were found using the *space-filling curves* approach described in [32]; protocols *d*_10_ and *d*_12_ were found using Sobol sensitivities [33] of the Wang and Beattie models, respectively; protocols *d*_4_ and *d*_5_ were found using a brute-force approach to maximise the sensitivity of model output to changes in parameters for the Beattie and Wang models, respectively; protocol *d*_7_ was found by considering 3-step blocks, randomising the durations of each step and optimising the voltages; whereas, protocol *d*_11_ was found by instead randomising the voltages and optimising the durations.

The remaining three protocols (*d*_1_, *d*_2_ and *d*_6_) were designed manually, without the use of an algorithm. Lei *et al*.’s *staircase* protocol [12], *d*_1_ was shown to permit the estimation of transition-rate parameters in models of I_Kr_. A similar protocol, *d*_2_ is included because it includes a central section in which there are many short-duration segments. We expect that this protocol highlights more of I_Kr_’s short-timescale behaviour (namely its *inactivation*/*recovery-from-inactivation* process, which occurs very rapidly). Finally, *d*_6_ is was performed without the intention of providing useful parameter estimates—in fact, our models are *practically unidentifiable* under *d*_6_ [10]. Nevertheless, this protocol consists of a sequence of action-potential voltage traces and, hence, provides physiologically relevant data for the purpose of model validation.

The protocols described above were performed sequentially before and after the addition of dofetilide, a drug known to specifically block I_Kr_ [34]. For QC, we perform four repeats of the *staircase* protocol, *d*_1_, allowing us to discard data from wells that do not remain stable over the course of the experiment. In particular, the *staircase* protocol, *d*_1_, was repeated four times, twice at the beginning of the experiment and twice after all other protocols. All other protocols were performed exactly once before, and once after, the addition of dofetilide. This procedure is illustrated by the schematic in Figure 4.

Each protocol contains some common elements which are included to aid postprocessing, as described in the following section. In particular, each protocol includes an identical section at the beginning of the protocol (the *leak* ramp). These sections allow for the estimation of leak-model parameters, and to infer the *reversal potential, E*_Kr_. Each protocol beings at the *holding potential, V*_cmd_(0) = −80mV, before *V*_cmd_ steps down to −120mV before gradually increasing back to −80mV. Because I_Kr_ is small in this range of voltages ([−120, −80]), this allows the determination of the leak current (Equation 7). A subsequent segment where *V*_cmd_ is held at +40mV permits validation of this leak model by checking that *I*_Kr_ = *I*_obs_ − *I*_L_ is positive during this step (as we would expect [9]). Similarly, a *reversal ramp* section is included at the end of each protocol, the apparent reversal potential of the channel, (that is, *E*_Kr_ in Equation 2) to be observed. This section begins with a +40mV preconditioning step, before *V*_cmd_ is rapidly reduced from −70mV to −110mV. The postprocessing methods applied to the leak-ramp and reversal-ramp sections of our protocols are described in the following section.

## 5 Fitting mathematical models to patch-clamp data

### 5.1 Postprocessing

#### Leak-model fitting

The leak ramp at the beginning of each voltage protocol, and the reversal ramp at the end of each voltage protocol, aids our postprocessing. We use the leak ramp to fit a linear leak-current model to the data, allowing us to subtract away leak current and observe the remaining current (which is dominated by I_Kr_). Details of this *leak-correction* procedure are provided in the Electronic Supplement (Appendix A). As mentioned in Section 3, we also perform *drug subtraction*, whereby our protocols are repeated after the addition of dofetilide, a known I_Kr_ blocker (after which we assume the maximal conductance, *g* = 0 and so *I*_Kr_ = 0. This is done as to minimise any endogenous (that is, non-I_Kr_) currents present in our postprocessed traces.

The results of our leak-correction, drug subtraction and reversal potential inference are used for quality control (QC). Our QC criteria largely follow those used in [26], which are, where possible, applied to all protocols. Though, we also include QC criteria involving *E*_obs_ and the relative sizes of the post-drug leak-corrected trace and the pre-drug leak-corrected traces. Full details regarding our postprocessing and QC procedures are provided in the Electronic Supplement (Appendix A).

#### Reversal potential inference

We infer *E*_Kr_ from the data using the reversal-ramp segment [26] at the end of each protocol. This is done using polynomial interpolation to find the point at which the current becomes negative (or, in other words, reverses). Following leak correction and drug subtraction, we estimate the reversal potential by fitting an order-4 polynomial to the current and use this to identify the time, *t\** at which *I*_Kr_(*t\**) = 0. We let *E*_obs_ = *V*_cmd_(*t\**) denote the *observed* reversal potential. To account for discrepancy between *E*_Nernst_ and *E*_obs_, we assume that any difference in these values is due to some voltage offset, that is, *V*_off_ = *E*_Nernst_ − *E*_obs_. This discrepancy and some alternative approaches are discussed in the Electronic Supplement (Appendix D).

### 5.2 Computational methods for model fitting

#### Forward simulation of Markov models

As introduced in Section 2, each of the four Markov models we employ may be seen as systems of ODEs. When *V*_m_ is held constant, our governing equation is a linear system of ODEs, for which there exists a range of computational methods [35]. However, the same is not true during the leak ramp and reversal ramp, where the solution cannot be expressed as a matrix exponential, and we instead resort to numerical integration methods. In particular, we use LSODA [36], an algorithm designed for the solution of stiff ODE systems. Here, we set both the relative and absolute tolerances to 10^*-*8^ to ensure the accuracy of our solutions.

#### Optimisation

We fit our models to time-series data by computing *maximum-likelihood estimates* (MLEs) under the assumption that our observations are subject to independently and identically distributed, additive Gaussian noise. That is, for a given protocol, *d*, we compute,

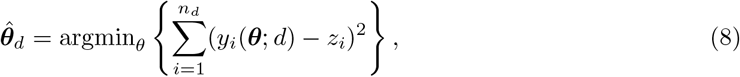

where *n*_*d*_ is the number of observations in protocol *d*, our *i*^th^ observation is denoted by *z*_*i*_, and the model output (for the *i*^th^ observation) for a given parameter vector ***θ*** is denoted by *y*_*i*_(***θ***; *d*). To fit our models, we seek a solution to this optimisation problem (Equation (8)). A general closed-form solution is not available, so we must resort to numerical optimisation methods.

To fit our models, we use CMA-ES [37], a stochastic optimisation method which proposes improved parameter estimates according to some continually updated sampling distribution, providing a chance to escape from local optima. Moreover, we repeat our optimisation 30 times from different initial sampling distributions, which allows us to explore more of the model’s parameter space and, provided we reliably recover the same parameter estimate, demonstrates that we are able to identify the true global optimum (that is, we can reliably compute Equation 8). For all models except the Wang model, we select the population size (that is, the number of parameter vectors sampled for each generation) by computing the integer,

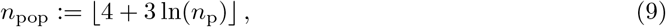

where *n*_p_ is the number of model parameters, using the heuristic suggested by the PINTS package [38] In the case of the Wang model, we instead increase the population size to 50. Open source code is publicly available, please see ‘Data Availability’ at the end of the manuscript.

#### Parameter space boundaries

We constrain our parameter space to mitigate the stiffness of our model’s governing equation (Equation 1). In particular, in line with [11], we ensure that no transition rate exceeds 10^3^ ms for at least one voltage in the range [−120 mV, +60 mV]. We also apply rather lenient bounds on individual model parameters, including the maximal conductance, *g*. Following [22] we perform each optimisation multiple times using different initial guesses. Here, we use 30 initial guesses for each sweep of each protocol. Our initial guesses for our ‘*a*’ and ‘*b*’ parameters are sampled from our parameter space using log-uniform and uniform distributions, respectively. Further details regarding these constraints and the sampling of initial guesses provided in the Electronic Supplement (Appendix C).

### 5.3 Results of model fitting and validation

#### Optimisation results

In order to be confident that we have successfully identified the optimal parameter set, we should expect that we obtain similar results from repeated runs of our optimisation procedure (which both starts from a randomised initial guess, and is inherently stochastic). Figure 5 shows the results of one particular optimisation task with 30 repeats. Here, we can see that amongst our best runs, the resulting parameter set varies only slightly—those results which correspond to a less than a 1% increase in RMSE when compared to the best found parameter set. In this case, these parameter sets occupy lie within a small region of parameter space, meaning that our optimisation methods are able to reliably find the correct parameter set.

**Figure 5:**
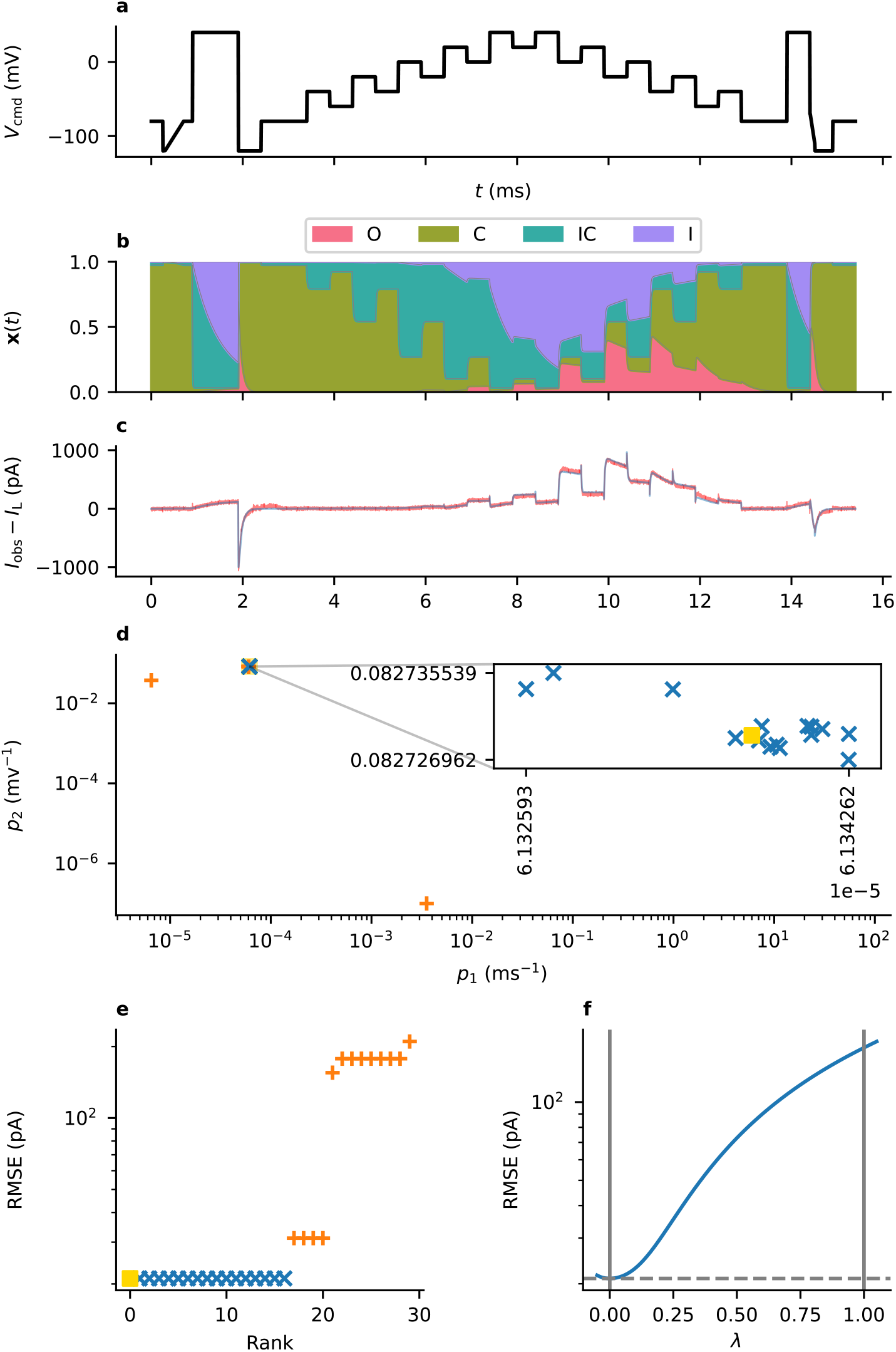
Results from fitting data from Well B20 with the Beattie model. Panel **a** shows the data and the model fit. Panel b: shows the membrane voltage during this protocol under the assumption that(*V*_cmd_ = *V*_m_). Panel **c** shows estimates of *p*_1_ and *p*_2_ obtained from thirty repeats of the optimisation routine—the blue markers shown in the inset correspond to those results where the RMSE is at most 101% of the minimum value. Panel **d** shows the amount of RMSE error in each fit. Panel **e** shows a cross-section through the likelihood surface, starting at our best estimate of the parameters (*A* = 0, yellow square) and finishing (when *A* = 1) at a parameter set with identical maximal conductance (*g*), but where the transition-rate parameters are taken from the model’s original publication (Cell # 5) [9].

#### Quantifying predictive accuracy

Not only do our models provide a good fit to the data (as exemplified in Figure 5), but the same parameter estimates perform well when applied to other protocols. This is largely true across each of our model structures, as shown in Figure 6, where we show the variability of these predictions—across our set of model structures and across wells.

**Figure 6:**
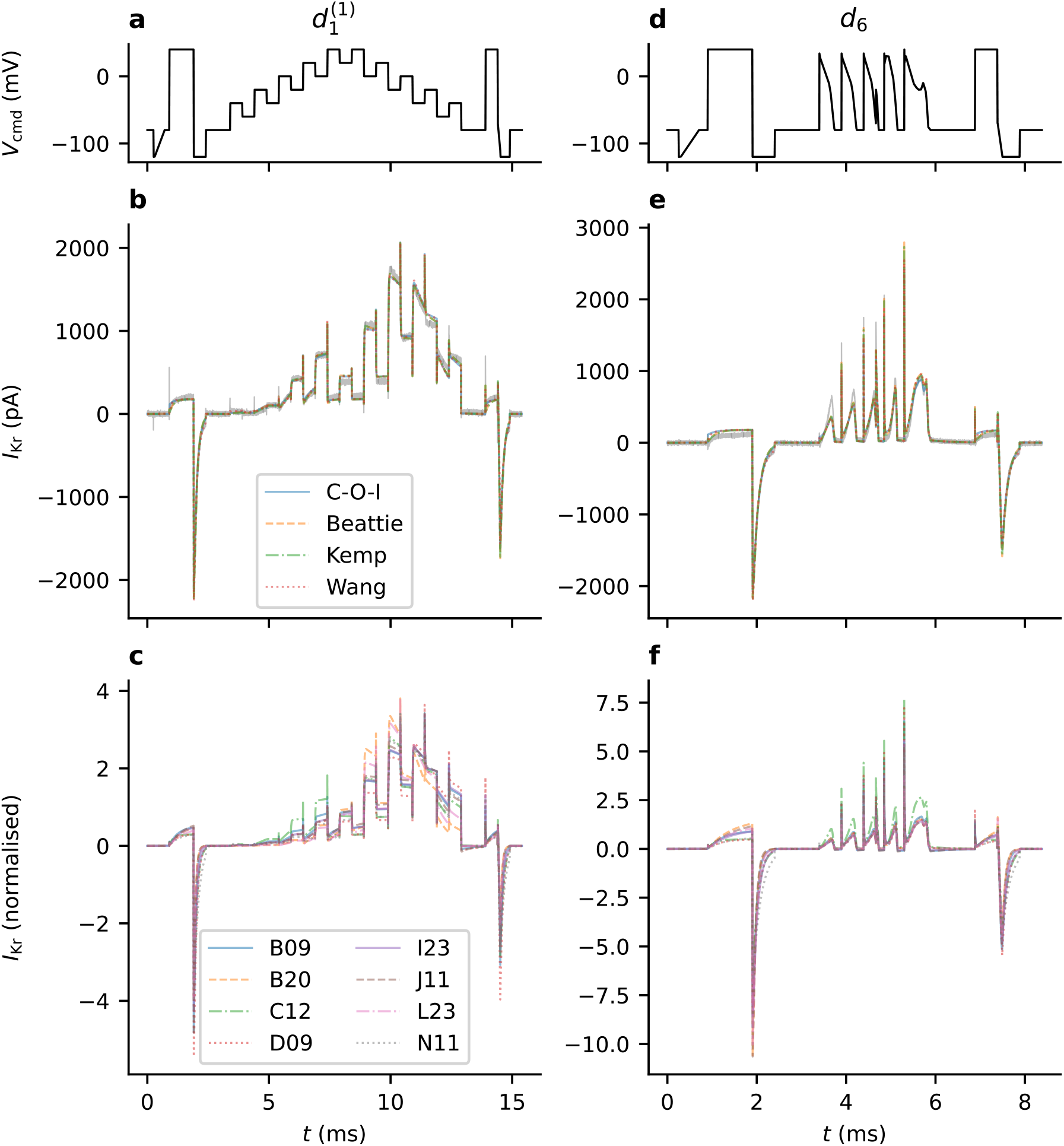
Our model predictions compared across model structures and across wells. All panels on the left relate to sweep 1 of the *staircase* protocol, 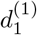: (**a**) the *d*_1_ protocol; (**b**) fits obtained using each of our models for one particular well, C12; (**c**) and fits obtained using one particular model (Beattie) [9] for each well. Similarly, all panels on the right relate to our validation protocol *d*_6_: (**d**) the *d*_6_ protocol; (**e**) predictions made using parameter estimates obtained from the fits shown in panel b, using each model, for the same well C12; and (**f**) Beattie model predictions of *d*_6_ using parameter sets obtained from multiple wells (fits shown in panel c). The model output shown in panels **c** and **f** is normalised such that the root-mean-square of each trace is 1.

From each of our parameter sets (each obtained from a different sweep of the data), we obtain an ensemble of parameter estimates. When input into our mathematical models, these parameter estimates give rise to an ensemble of predictions. In Figure 7, we show which sections of our voltage protocols prove particularly difficult to fit by computed a weighted average of the residuals of our fitted models at each time point. This average is weighted according to the size of our noise estimate, that is, we compute, 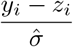, and average this quantity across wells. The resultant figure, Figure 7, therefore, highlights the sections of our protocols for which the Beattie model consistently over- or under-estimates I_Kr_. Similar figures are shown for the other models in the Electronic Supplement (Appendix E).

**Figure 7:**
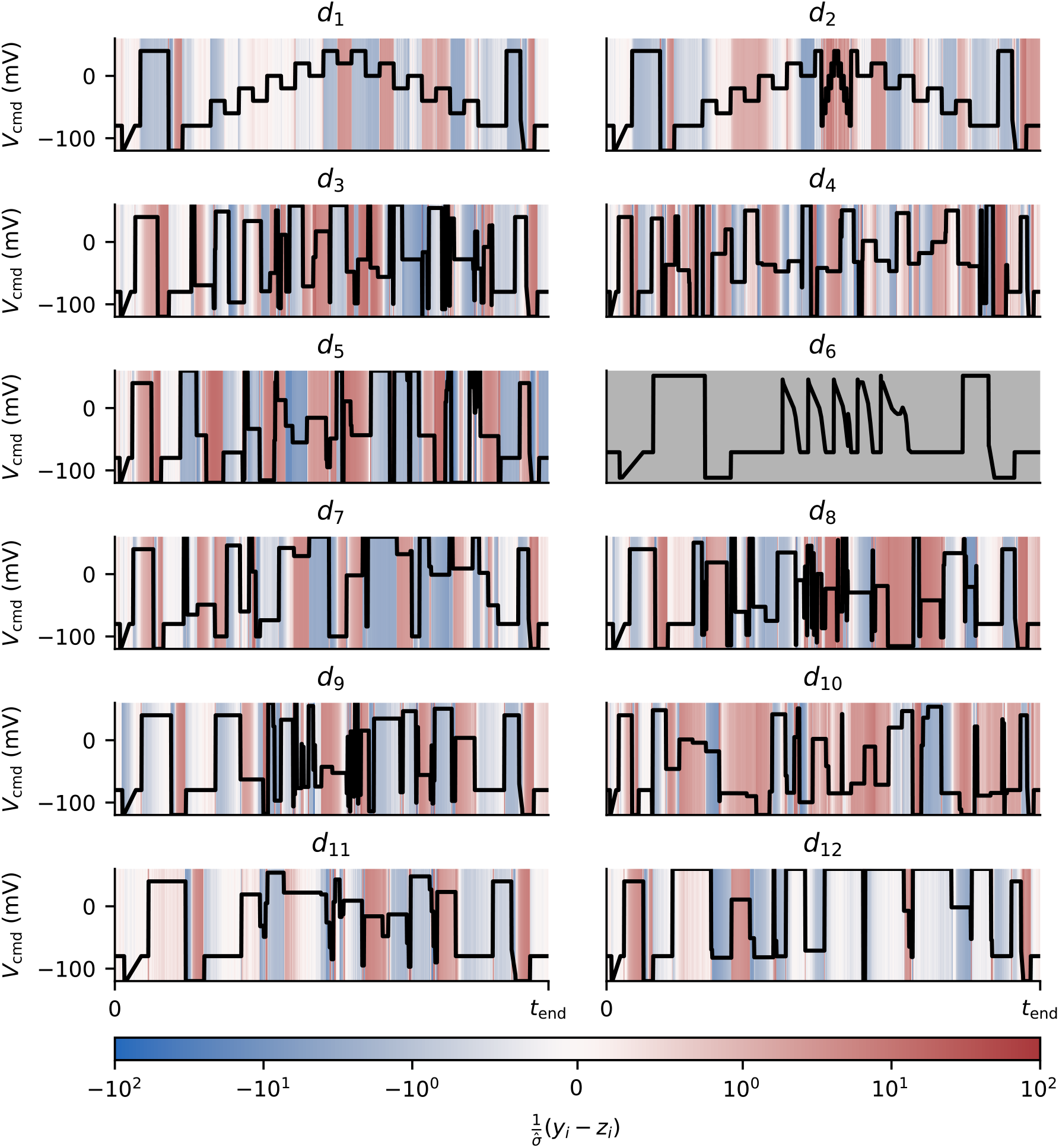
Residuals obtained when fitting the Beattie model to each training protocol (averaged across wells). The values plotted for 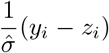 have been clipped to lie between -100 and +100. Protocol *d*_6_, which is used only for validation and not fitting, is shown in grey.

Likewise, Figure 8 shows sections of our protocols where the Beattie model (when fit to using a range of protocols) consistently over- or under-predicts I_Kr_. Here, we quantify the tendency of a model’s predictions to be consistently discrepant at each time-point, *t*_*i*_, by calculating,

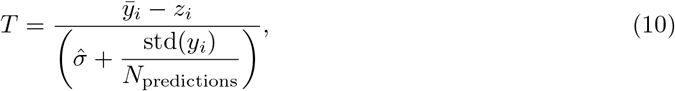

where 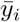 and std(*y*_*i*_) are the mean and standard deviation, respectively, of our *N*_prediction_ model predictions at time *t*_*i*_. When *T* ≫ 0 or *T* ≪ 0, this statistic shows that our ensemble of predictions consistently makes inaccurate predictions. Moreover, the sign of *T* indicates where models tend to over- or under-estimate I_Kr_. Here, we only include predictions and not model fits—that is, we discard parameter estimates obtained from the protocol under consideration. Plots of this type are shown for each model structure in the Electronic Supplement (Appendix E).

**Figure 8:**
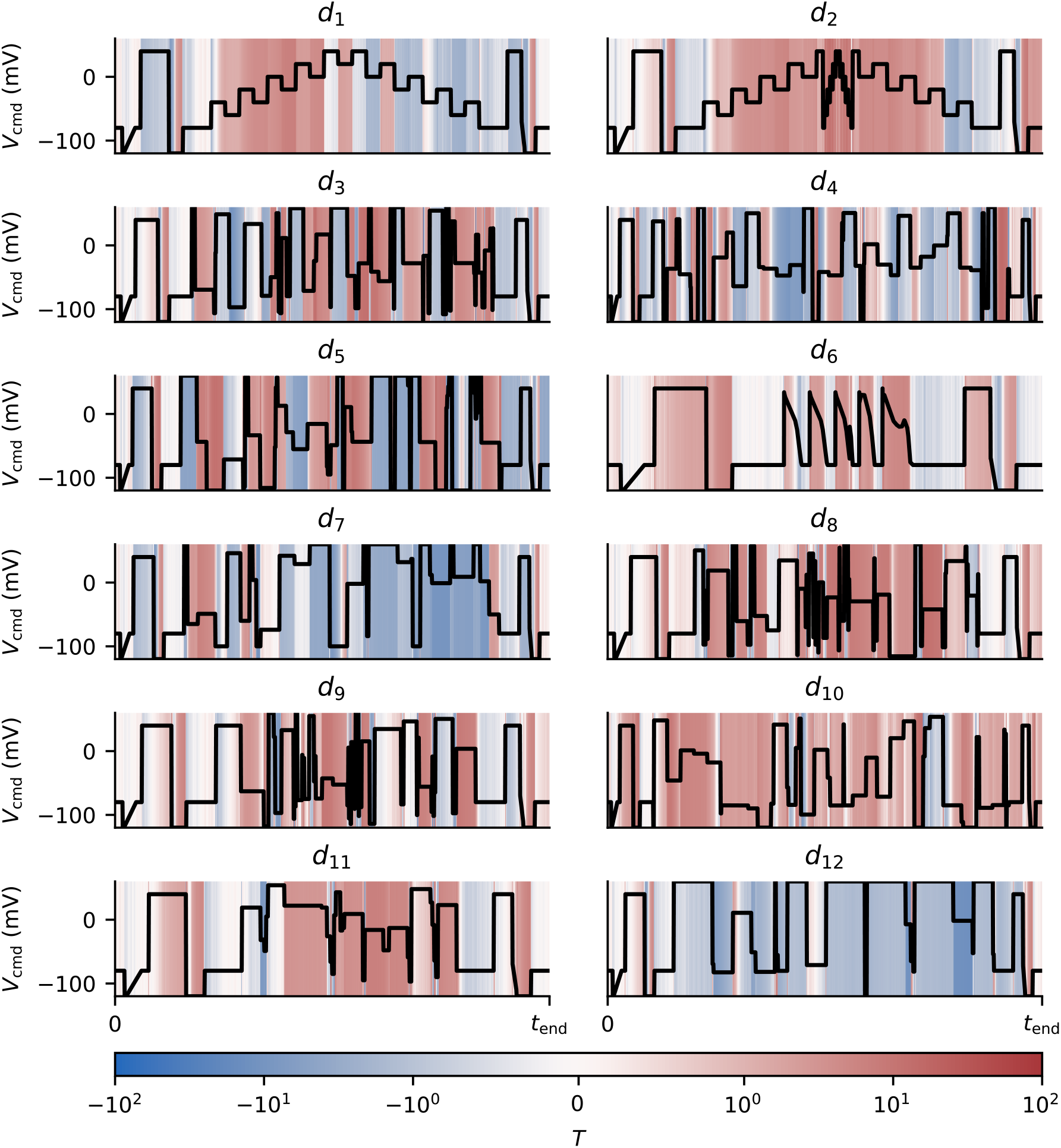
The average behaviour of the Beattie model when producing predictions for unseen protocols, averaged across wells. Sections of the protocols highlighted in red show where the model consistently overestimates I_Kr_, as quantified by our *T* statistic (Equation (10)). Whereas, sections highlighted in blue show consistent underestimation. The *T* values are clipped between −100 and +100.

For the purpose of comparing predictive accuracy across models, we quantify the accuracy of model predictions, using the *normalised* root-mean-square error,

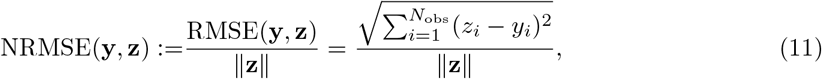

where RMSE denotes the (non-normalised) root-mean-square error. As the maximal conductances of different cells may vary, this is a way to fairly compare and average predictive performance across wells. It also allows us to fairly compare protocols with different numbers of observations.

The accuracy of these model predictions (indexed by the sweeps used for fitting and those used for validation) may be summarised in a heatmap, as shown in Figure 9. This shows the accuracy of each member of our ensemble of predictions two particular wells (Well D09 and Well B20), and a particular choice of model structure (the Beattie model [9]). Then, we take the average of these values and show (for each candidate model structure) the average predictive accuracy under each pair of fitting and validation protocols. Figure 10 compares each model’s averaged, cross-validation heatmap.

**Figure 9:**
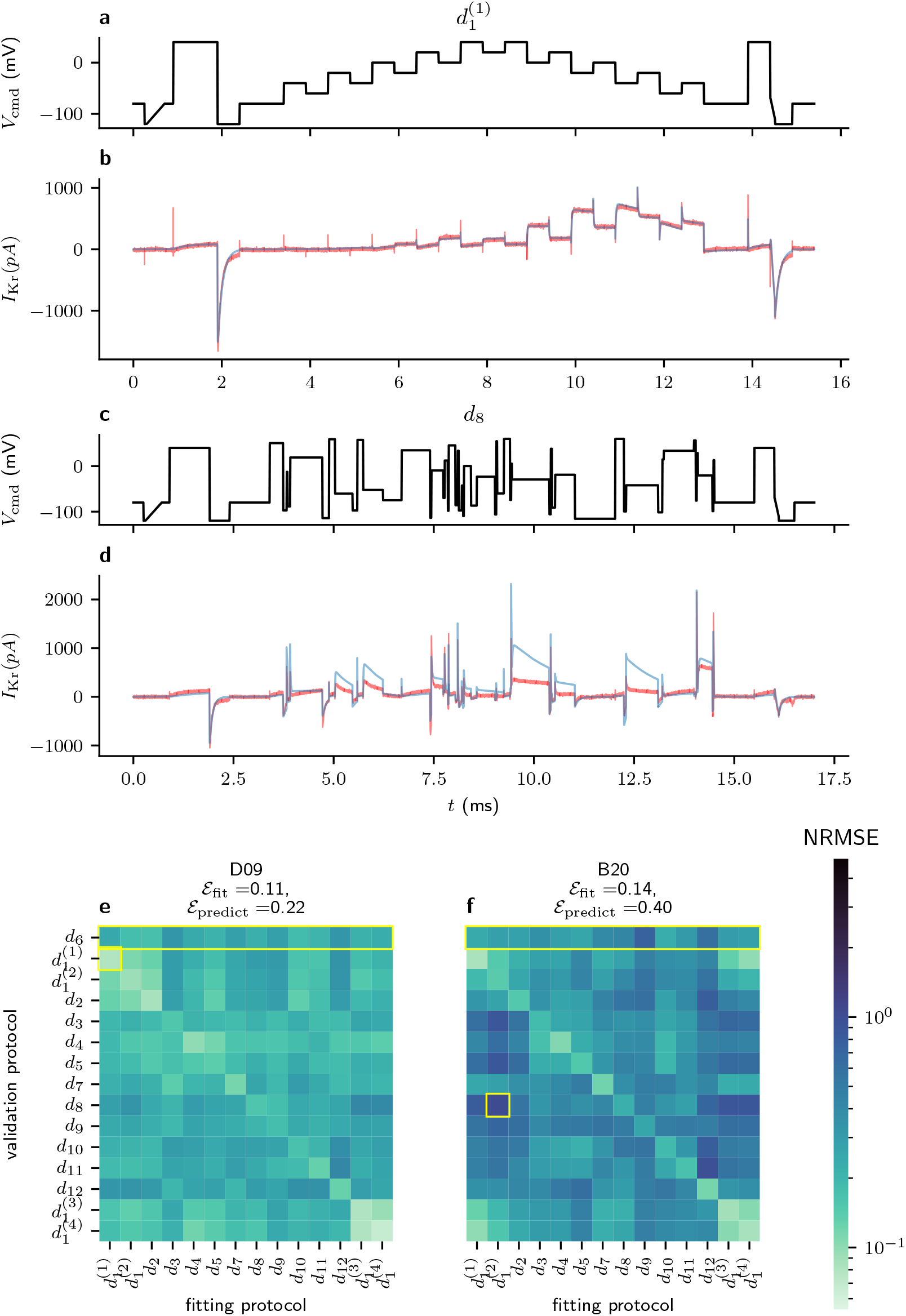
A cross-validation heatmap showing the predictive accuracy of the Beattie model. The two wells shown, Well D09 (panel **e**) and Well B20 (panel **f**), are the wells with the greatest and least average NRMSE across each pair of fitting and training sweeps. Panel **b** shows the worst model prediction (in terms of the NRMSE) from Well D09, corresponding to the highlighted cell in panel **e**. For comparison, Panel **c** shows the lowest scoring (NRMSE) cell in panel **f**. The corresponding voltage traces are shown in panels **a** and **c**.

**Figure 10:**
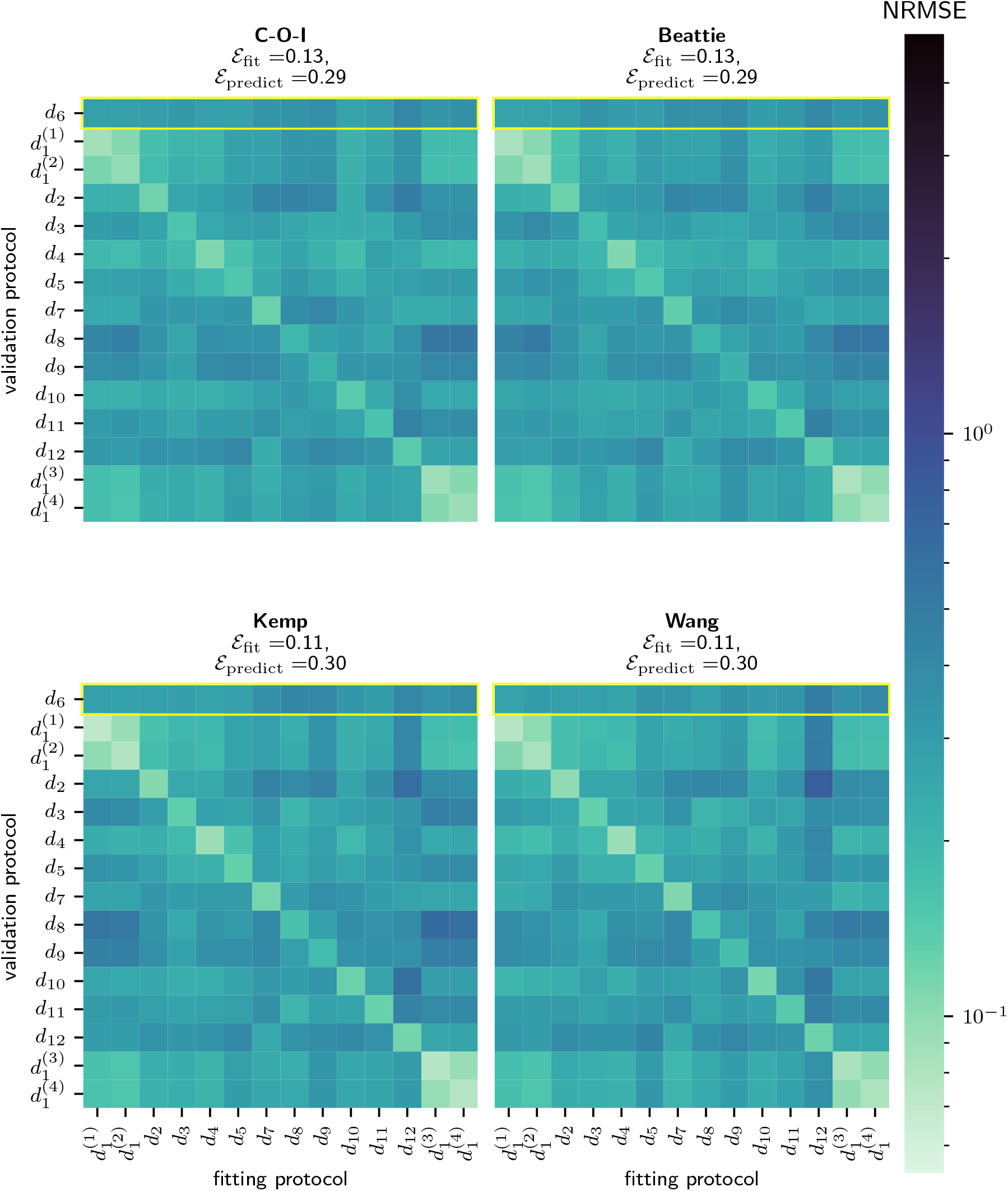
A comparison of the predictive performance of our chosen model structures. Each heatmap shows the average normalised root-mean-square error when the given model is trained and validated each pair of protocols.

## 6 Variability in parameter estimates

By fitting our data to each sweep in our dataset, we obtain a collection of parameter estimates for each model structure, where each individual parameter-estimate vector, 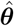, pertains to a particular sweep in our dataset. For each Markov model, and for each of the 8 wells selected by QC, there are 14 parameter estimates—one parameter estimate arising from each repeat of each fitting protocol (a single repeat of each protocol except *d*_6_, and an additional three repeats of *d*_1_). In this section, we discuss the variability of these parameter estimates using a simple, linear statistical model.

Plots of our parameter estimates, shown in Figure 11, suggest a relationship between the well and protocol an estimate was obtained from and its value; some protocols yield consistently high estimates of *p*_1_ when compared to other protocols, for example. Since these plots (Figure 11) only show two parameters in particular, *p*_1_ and *p*_2_, which determine a single transition rate, *k*_1_ = *p*_1_ exp{*p*_2_*V* } in the Beattie model [9].

**Figure 11:**
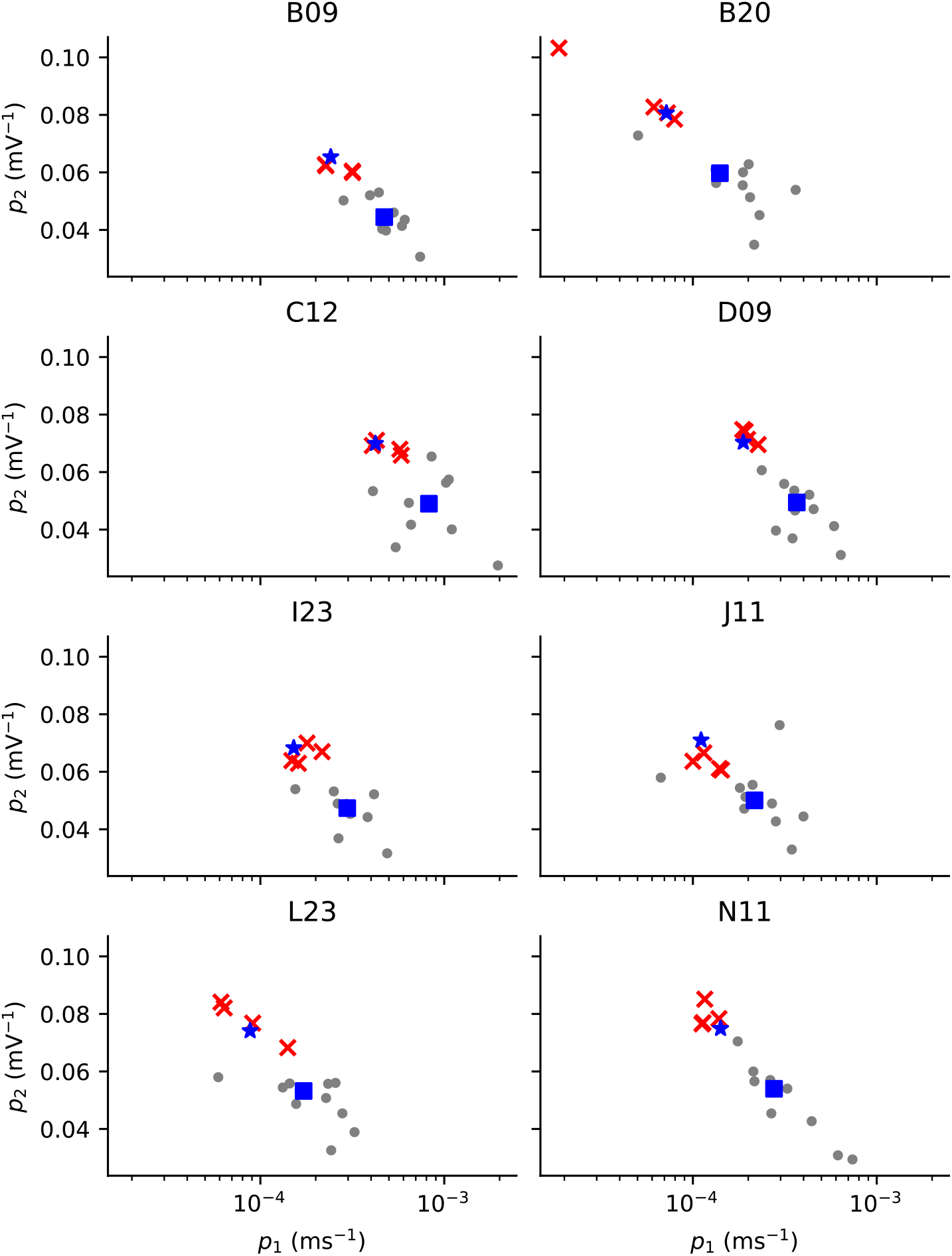
A linear statistical regression model, ℳ_w,d_, largely fits and recapitulates the well- and protocol-dependence in mechanistic model parameter estimates for the Beattie model. Each panel shows the same two parameter estimates obtained from a given well, and highlights the parameter estimates obtained from the *staircase* protocol (red crosses), note that these are just two of the eight model kinetics parameters shown as an illustration, but the linear model is constructed for all eight. The blue square shows the well-dependent effect according to ℳ_w_, our linear model with no protocol-dependent effects, and the blue star shows the sum of the well and protocol effects (for the given well and the *d*_1_ *staircase* protocol) according to the ℳ_w,d_ model. Note that *p*_1_ (the *abscissae*) has been log-scaled, as it is in our linear model.

We fit a simple statistical linear regression model ℳ_w,d_ for our parameter estimates, including well (w) and protocol/design (d) effects. Where our models contain transition rates of the form, *k* = *A* exp {±*bV*} = exp {*a* + *bV*}, we fit our linear model using *A* and *b*. Only parameters related to the transition-rate matrix **Q** are included in this analysis because we anticipate significant cell-cell variability in maximal conductances, whereas we expect transition rates to be closely related to biophysical constants [10].

To test whether there is significant well- and/or protocol-dependence in our parameter estimates, we use log-likelihood differences (LLDs). We do this by considering,

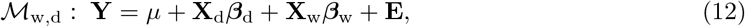

where: **Y** is an *N*_trace_ × *N*_p_ matrix where *N*_trace_ is the number of fitting traces and *N*_p_ is the number of parameters in our I_Kr_ model, such that each row is a parameter estimate obtained from model fitting; **X**_w_ and **X**_d_ are well- and protocol-effect *design matrices* where each row encodes the well/protocol used for a given parameter estimate; *β*_w_ and *β*_d_ are our parameter matrices, with each row representing a particular well or protocol effect (respectively), and with each column corresponding to a different parameter in our I_Kr_ model; **E** is a matrix of random errors, such that for each *k* ∈ {1,…, *N*_p_}, the errors in the *k* column are IID Gaussian distribution random variables with expectation zero and unknown standard deviation *σ*_*k*_. So that this model is identifiable, we insist that the protocol and well effects sum to zero, that is, 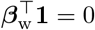 and 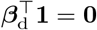. The remaining models, ℳ_w_, ℳ_d_, and ℳ_0_ are set by the constraints, *β*_w_ = 0, *β*_d_ = 0, and *β*_d_ = *β*_w_ = 0, respectively.

We list the maximum likelihood of each model, ℳ_*i*_, for each set of parameter estimates (arising from our candidate I_Kr_ model structures) in Table 1, as well as the resulting log-likelihood differences between models ℳ_w,d_ and ℳ_d_ (**LLD-W**), and ℳ_w,d_ and ℳ_w_ (**LLD-D**). Thus, **LLD-W** is a statistic quantifying the significance of the well-dependent effects, and **LLD-D** is a statistic quantifying the significance of the protocol-dependent effects. From the values presented in Table 1, we can see that including the well-dependent and protocol-dependent effects leads to a large increase in likelihood. This indicates that there is significant protocol- and well-dependence in our parameter estimates. The parameter estimates that we obtained are shown in Figure 11. From these results, we can see that there is noticeable variability between parameters obtained from different wells under the same protocol, and also variability between the estimates taken from the same protocol, but from different wells.

**Table 1:** Log-likelihoods and likelihood ratios (log-likelihood differences, LLD) for each of the linear regression models, applied to all of our biophysical models (that is, the collection of parameter estimates obtained using each biophysical model). Here we see that in each case, and for each biophysical model, that both well- and protocol-effects are very significant. Whilst **LLD-D** suggests that there is discrepancy between the recordings of I_Kr_ (taken from each well) and the dynamics of our biophysical models, the magnitude of **LLD-W** (across all model structures) suggests the presence of latent, well-dependent effects.

| Model | $\mathcal{M}_0$ | $\mathcal{M}_w$ | $\mathcal{M}_d$ | $\mathcal{M}_{w,d}$ | LLD-W | LLD-D |
| --- | --- | --- | --- | --- | --- | --- |
| C-O-I | 1465.6 | 1668.5 | 1724.4 | 2111.1 | 386.7 | 442.6 |
| Beattie | 1563.3 | 1817.0 | 1784.2 | 2247.2 | 462.9 | 430.2 |
| Kemp | 931.8 | 1099.8 | 1113.7 | 1356.3 | 242.6 | 256.5 |
| Wang | -2082.1 | -1976.1 | -1836.7 | -1673.5 | 163.2 | 302.6 |

## 7 Discussion

We have collected new data using multiple information-rich experimental designs and used these to thoroughly validate our models of I_Kr_. The methods used to process this data, including our fully-automated Quality Control (QC) procedure, are suitable for future work involving the collection and analysis of multiprotocol patch-clamp experiments for I_Kr_, and possibly other ion-channel currents. Note that whilst we observed a low success rate (according to our strict QC criteria, as detailed in Appendix B of the Electronic Supplement), the data retained after QC is of high quality and shows remarkable fidelity with our mathematical models. Even so, future improvements to the experimental equipment and methodology may result in an increased success rate and, perhaps, even cleaner data.

Because our data were collected from a diverse range of information-rich experiments, we are able to thoroughly validate the predictive accuracy of a small selection of I_Kr_ models. Overall, we have shown that each model is able to accurately recapitulate our I_Kr_ recordings. This is particularly true in certain wells, such as Well D09, where our models not only provide very accurate fits to the data, but are able to predict the current during unseen protocols to a high degree of accuracy (see Figure 10). Broadly speaking, models with more parameters (the Kemp [29] and Wang [28] models) produced more accurate fits to our data, as quantified by *ε*_train_, whereas the simpler models (the C-O-I and Beattie [9] models) produced slightly more accurate model predictions for unseen protocols (see Figure 10).

However, the difference between our competing model structures regarding *ε*_train_ and *ε*_pred_ are rather subtle. Perhaps our ideal patch assumptions (Equation 5 and 6) result in models too discrepant to allow such differences in model structure to make a material difference in these values. If these ideal-patch assumptions are relaxed (for example, by the inclusion of experimental artefacts [24]), the data herein should prove valuable for the training and validation of further I_Kr_ models. Our work is ongoing in this regard.

Our statistical analysis of our ensembles of parameter estimates suggest that there are strong well- and protocol-dependent effects acting upon our parameter estimates. As discussed in Shuttleworth *et al*. [22], we expect to see these protocol dependent effects when there is discrepancy between our mathematical models and the underlying biophysical mechanisms that we observe. The strong well-dependent effects, however, suggest substantial experimental variability, unaccounted for by the models presented here. As argued by Lei *et al*. [24], it is possible that well-dependent *experimental artefacts* are the dominant cause of this well-to-well variability. Perhaps the large well-to-well variability in our kinetic parameters is caused by the overfitting of kinetic parameters when these effects are omitted. Further work will investigate whether the inclusion of such artefact effects decreases this well-to-well variability, and improves accuracy of our model predictions. If so, the inclusion of artefact effect may result in yet more accurate predictive models of I_Kr_ and allow us to better discriminate between competing model structures on the basis of predictive accuracy and the protocol-dependence of parameter estimates.

## Data Access

Open source code for all the model fitting and plots in this paper can be found on https://github.com/CardiacModelling/multiprotocol_data_fitting.

## Acknowledgements

Model fitting was performed on The University of Nottingham’s Ada High-Performance Computing resource.

## Funding

This work was supported by the Wellcome Trust [grant no. 212203/Z/18/Z]; the Science and Technology Development Fund, Macao SAR (FDCT) [reference no. 0155/2023/RIA3 and 0048/2022/A]; the University of Macau [reference no. SRG2024-00014-FHS and FHS Startup Grant]; the EPSRC [grant no. EP/R014604/1]; and the Australian Research Council [grant no. DP190101758]. GRM acknowledges support from the Wellcome Trust via a Wellcome Trust Senior Research Fellowship to GRM. CLL acknowledges support from the FDCT and the University of Macau to CLL. We acknowledge Victor Chang Cardiac Research Institute Innovation Centre, funded by the NSW Government.

This research was funded in whole, or in part, by the Wellcome Trust [212203/Z/18/Z]. For the purpose of open access, the authors have applied a CC-BY public copyright licence to any Author Accepted Manuscript version arising from this submission.

## Notes

https://github.com/CardiacModelling/multiprotocol_data_fitting

### Competing Interest Statement

The authors have declared no competing interest.

