## Supplementary Material for "Evaluating the predictive accuracy of ion channel models using data from multiple experimental designs"

### A Postprocessing a multiprotocol dataset

As described in Section 5, we perform postprocessing prior to fitting our models. In particular, we perform leak correction, drug subtraction, to infer the reversal potential. These results are then used to perform quality control (QC). In this section, we explain these methods and supplement the main text with additional results.

**Leak correction** Our leak-current model (Equation 7 of the main text) quantifies the current flowing through the imperfect seal between the cell and the well-plate. This is an equivalent circuit formulation corresponding to a simple resistor. Following [12], we estimate our leak-model parameters (that is, $g_{\mathrm{L}}$ and $E_{\mathrm{L}}$) using the leak-ramp section of our protocols. We then use the linear-leak model to compute $I_{\mathrm{L}}$, and subtract the resulting current from our observed data. Under our ideal-patch assumption, this removes the leak current from our data, allowing us to calibrate our mathematical models.

Fitting the parameters of the leak model using the leak ramp is possible because $\mathrm{I_{Kr}}$ is typically small for the range of voltages used in the leak-ramp section. Hence, we may assume that our observed current is entirely due to the leak current that is,

$$I_{\mathrm{obs}}(t) = I_{\mathrm{L}}(t), \tag{A.1}$$

for the duration of the leak ramp.

Using simple linear regression, we write,

$$I_{\mathrm{obs}}(t_i) = \beta_1 + V_{\mathrm{cmd}}(t_i)\beta_2, \tag{A.2}$$

where our dependent variable, $I_{\mathrm{obs}}(t_i)$, is the raw data observed during the experiment at the $i^{\mathrm{th}}$ observation time, our independent variable, $V_{\mathrm{cmd}}(t_i)$, is the corresponding command voltage, and $\beta_0$ and $\beta_1$ are our regressor variables. These variables are related to our leak-model parameters by noting that $\beta_1 = -g_{\mathrm{L}}E_{\mathrm{L}}$ and $\beta_2 = g_{\mathrm{L}}$.

It is possible to perform this simple linear regression using only two distinct $V_{\mathrm{cmd}}$ values, thus a "leak step" may be used in place of our "leak ramp". However, the use of the leak ramp produces data for a range of voltages, allowing us to ensure the validity of the leak model. The +40mV step immediately following the leak ramp is also useful for validation because at the onset of this ramp, $\mathrm{I_{Kr}}$ activates slowly. This means that $I_{\mathrm{Kr}}$ remains small for some time, during which the observed current, $I_{\mathrm{obs}}$, mostly consists of $I_{\mathrm{L}}$. Hence, the observation of a significantly negative current here would indicate that the leak has been over-subtracted. The estimation of our leak-model parameters is shown in Figure A.1.

**Drug subtraction** Although our leak-correction method allows us to infer and correct for linear leak currents, there may be biological currents (other than $\mathrm{I_{Kr}}$ that are active in our cells. These currents are called *endogenous* currents, and we wish to minimise their presence as to avoid the pollution of our data. As mentioned in Section 3, we apply dofetilide to our cells and repeat each of our protocols after $\mathrm{I_{Kr}}$ has been completely blocked. This results in two sets of traces: pre-drug traces which are recorded before the addition of any drug, and post-drug traces which are recorded after $\mathrm{I_{Kr}}$ has been completely blocked [9]. Since a leak current is present in both cases, we fit leak-current parameters to each trace. This is done independently for each trace to account for changes in the leak current over the course of the experiment—it is plausible that the quality of the seal would degrade over time (that is, $g_{\mathrm{L}}$ may increase over the course of the experiment).

By subtracting the leak-corrected post-drug traces from the leak-corrected pre-drug traces, we (ideally) eliminate all biological, non-$\mathrm{I_{Kr}}$ currents, so that we can fit our $\mathrm{I_{Kr}}$ models directly to our postprocessed data. To compute our postprocessed trace (that is, the current remaining after leak correction and drug subtraction), which we use for model fitting, $I_{\mathrm{post}}$, we subtract the leak-corrected

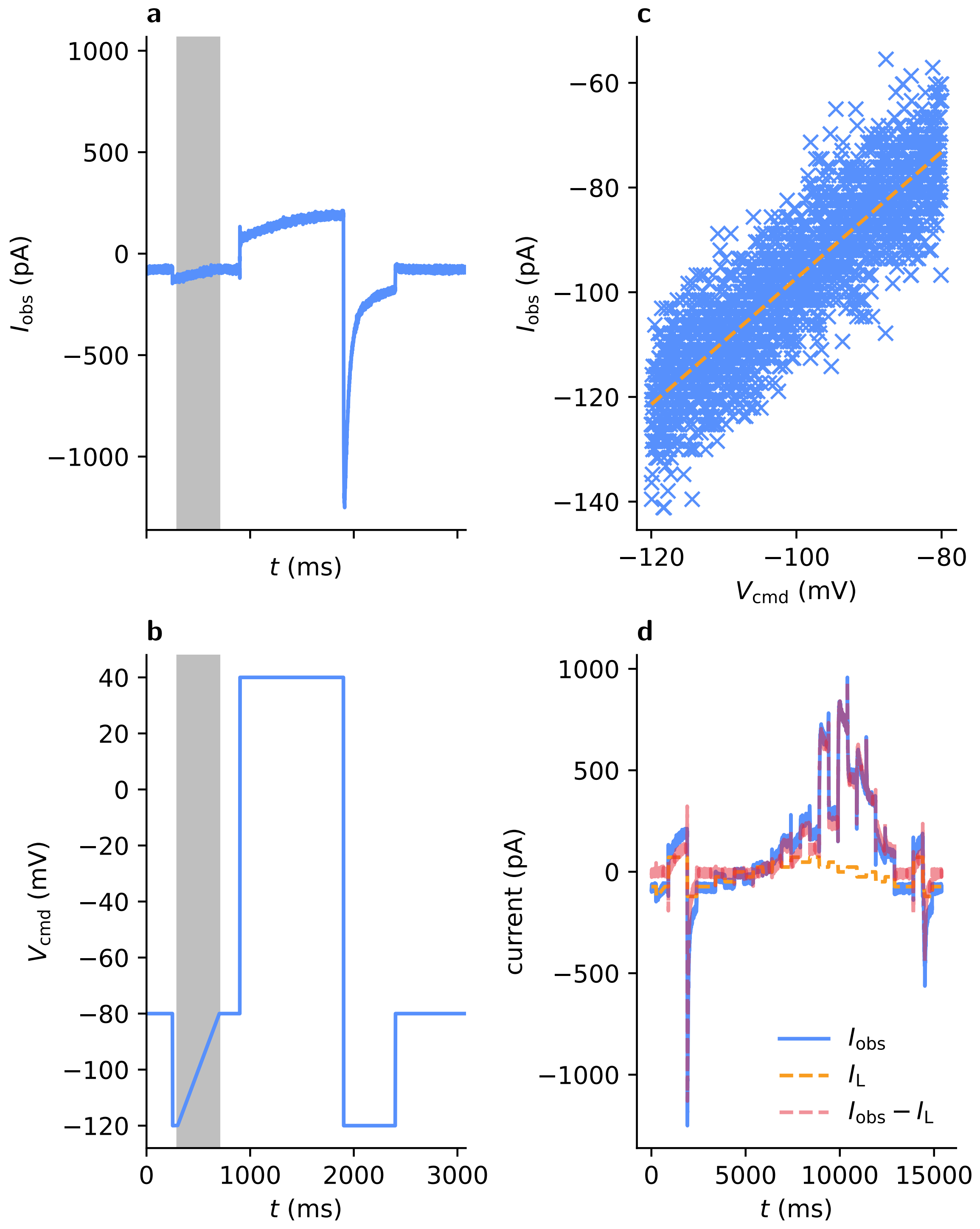


Figure A.1: Our linear-leak current model is fitted using simple linear regression. Panels **a** (showing the observed current) and **b** (showing the command voltage) highlight the leak ramp. Panel **c** shows the simple linear regression used to fit the linear leak model. Panel **d** shows the observed current ($I_{\text{obs}}$), the leak current ($I_{\text{L}}$) and the leak-corrected current ($I_{\text{obs}} - I_{\text{L}}$). This data was taken from the application of the *staircase* protocol to well B20.

pre-drug trace, $I_{\text{obs}}^{(\text{after})}$, from the leak-corrected post-drug trace $I_{\text{obs}}^{(\text{before})}$,

$$I_{\text{post}} = \left(I_{\text{obs}}^{(\text{before})} - I_{\text{L}}^{(\text{before})}\right) - \left(I_{\text{obs}}^{(\text{after})} - I_{\text{L}}^{(\text{after})}\right). \tag{A.3}$$

We may quantify the amount of nuisance, leftover current by considering the relative size of our leak-corrected traces before and after the addition of dofetilide,

$$R_{\text{leftover}} = \frac{\|I_{\text{obs}}^{(\text{after})} - I_{\text{L}}^{(\text{after})}\|}{\|I_{\text{obs}}^{(\text{before})} - I_{\text{L}}^{(\text{before})}\|}. \tag{A.4}$$

Under our ideal patch-clamp assumptions, we expect that $R_{\text{leftover}}$ is small (though non-zero due to the presence of random noise). This statistic provides a useful quantification of the relative size of current that remains after drug subtraction and leak correction. This leftover current may be due to endogenous currents, nonlinear leak, or miscalibration of the leak model (perhaps due to data pollution, or changes in the quality of the seal over time). This statistic, $R_{\text{leftover}}$, is also used for quality control as explained in the following section. An example of leak-correction and drug subtraction is shown in Figure A.2. Here, two consecutive sweeps of the *staircase* protocol, $d_1$, are shown alongside corresponding post-drug traces.

**Reversal potential inference** We do this by performing polynomial interpolation on the command voltage and leak-corrected current during the reversal ramp, that is, we take $V_{\text{cmd}}$ to be the independent variable, and $I_{\text{corrected}}$ to be the dependent variable fit a fourth-order polynomial, denoted $P(V_{\text{cmd}})$ to the data: pairs of $(V_{\text{cmd}}(t), I_{\text{obs}}(t))$. We then find roots of this polynomial, that is, $x$ such that $P(x) = 0$ which lie inside the voltage-range used during the reversal ramp, that is, with $-110\text{mV} < x < -70\text{mV}$. Furthermore, we insist that only one such $x$ exists. If we find a $P(x)$ for which there is not a unique root, $x$, satisfying these conditions, we discard the sweep (We refer to this criterion as **QC.Erev**). If there is a unique root, $x^*$ of $P(x)$, the reversal potential is $x^*$ because, due to our ideal patch-clamp assumptions, $I_{\text{out}} - I_{\text{L}} = I_{\text{Kr}} = 0$ implies that $V_{\text{m}} - E_{\text{Kr}} = 0$ (since we know that $x_{\text{O}} \neq 0$).

**Details of voltage clamp protocols** A summary of the properties of these experimental protocols is provided in Table A.1. These voltage protocols are also shown in Figure 3 of the main text.

### B Quality control

**Using Lei *et al.*'s quality-control criteria [12]** Before fitting our models, we use a set of *quality control* (QC) criteria to filter out failed experiments from our dataset. This is done to remove data from failed experiments, or data exhibiting low levels of $\text{I}_{\text{Kr}}$ (in relation to the size of the observational noise), instability, *etc.*.

The Nanion SyncroPatch automatically applies a short protocol and infers a number of parameters before each of our voltage protocols ($d_1$–$d_{12}$). Namely, these parameters are the seal resistance, $R_{\text{seal}} = \frac{1}{g_{\text{L}}}$, the capacitance of the cell membrane $C_{\text{m}}$, and the *series resistance*, $R_{\text{series}}$ [26]. We use these *machine estimates* to discard unsuitable traces from our dataset. As in [12], we ensure that for each sweep that each of these values lies within a predetermined range of plausible values,

$$0.1\,\text{G}\Omega < R_{\text{seal}} < 10\,\text{G}\Omega, \tag{B.1}$$
$$1\,\text{M}\Omega < R_{\text{series}} < 25\,\text{M}\Omega, \tag{B.2}$$
$$\text{and } 1\,\text{pF} < C_{\text{m}} < 10\,\text{pF}. \tag{B.3}$$

This criterion is referred to as **QC1**, and the subcriteria as **QC1.Rseries**, **QC1.Rseal** and **QC1.Cm** (each corresponding to a different variable).

We then ensure that the machine estimates, obtained from consecutive sweeps of the *staircase* protocol are not too dissimilar, as this would suggest instability in the experimental setup. we refer

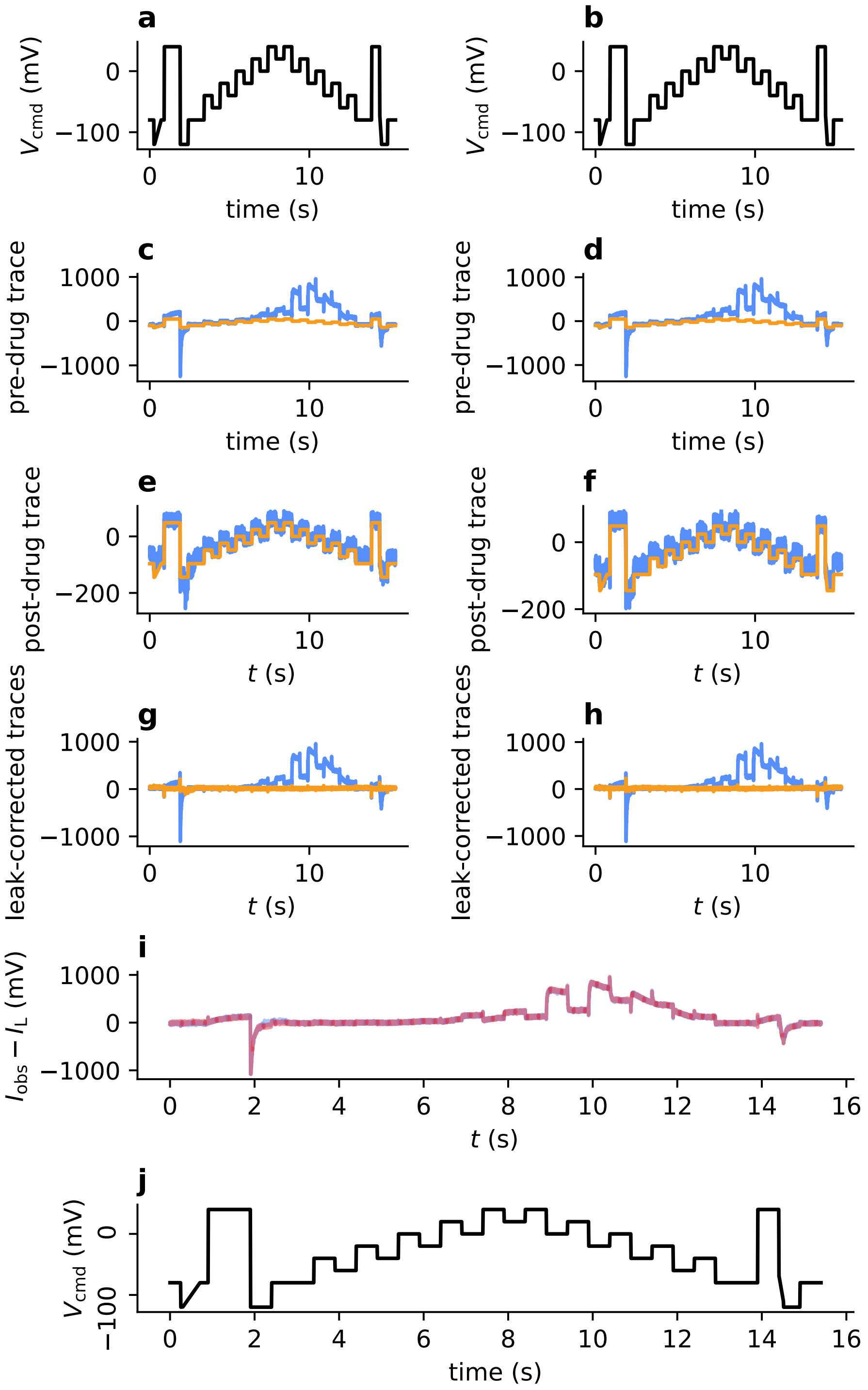


Figure A.2: Leak-correction and drug-subtraction performed as part of the postprocessing of our patch-clamp data. The figure shows two sweeps of $d_1$, with corresponding post-drug sweeps, which are subtracted from the pre-drug sweeps. Panels **a**, **b** and **j** show $V_{cmd}$ for the $d_1$ protocol. Panels **c** and **d** show the pre-drug traces (blue) and fitted leak currents (orange) for the first and second sweeps, respectively. Similarly, panels **e** and **f** show the post-drug traces and corresponding fitted leak currents. Panels **g** and **h** show the pre-drug (blue) and post-drug (orange) leak-corrected traces. Finally, panel **i** shows overlaid leak-corrected, drug-subtracted traces from the first (blue) and second (red) sweeps. These sweeps mostly overlap, showing excellent consistency over time.

| Protocol | duration (s) | no. segments | description |
|---|---|---|---|
| $d_1$ | 14.95 | 32 | *staircaseramp* |
| $d_2$ | 14.95 | 52 | *staircaseramp* with inverted and rapid centre |
| $d_3$ | 18.8 | 64 | a space-filling voltage protocol [32] |
| $d_4$ | 32.9 | 58 | found using brute-force sampling together with the Beattie model |
| $d_5$ | 22.5 | 58 | found using the brute-force approach with the Wang model |
| $d_6$ | 8.4 | 63 | action-potential protocol, used for validation |
| $d_7$ | 21.4 | 43 | found by maximising model differences under randomly chosen step durations |
| $d_8$ | 17.0 | 64 | a space-filling design [32] |
| $d_9$ | 14.6 | 64 | a space-filling design [32] |
| $d_{10}$ | 32.6 | 58 | found by maximising first-order Sobol sensitivities under the Wang model |
| $d_{11}$ | 21.1 | 43 | found by maximising model differences under randomly chosen voltage levels |
| $d_{12}$ | 22.4 | 40 | found by maximising first-order Sobol sensitivities under the Beattie model |

Table A.1: The duration of and number of segments (voltage steps or ramps) in each protocol that we performed. During these segments, the command voltage is either held constant, or changes according to some constant gradient (a ramp). Here, the number of segments includes segments which are common to all of our protocols—the leak ramp and reversal ramp, and their associated +40mV pre-conditioning steps.

to these criteria as **QC4.Rseal**, **QC4.Rseries** and **QC4.Cm**—each criteria again corresponding to a particular machine estimate. This is done by discarding wells for which,

$$\frac{\mathrm{std}(C_\mathrm{m}^{(2)}, C_\mathrm{m}^{(1)})}{\mathrm{mean}(C_\mathrm{m}^{(2)}, C_\mathrm{m}^{(1)})} < 0.25, \tag{B.4}$$

where $\mathrm{std}(\cdot,\cdot)$ and $\mathrm{mean}(\cdot,\cdot)$ denote the population standard deviation and sample mean, respectively, of the two values.

The remaining criteria (as in [26]) concern time-series data. Whilst they were originally applied to data obtained from Lei *et al.*'s *staircase* protocol, some may be directly applied to our other protocols, due to their shared features. Moreover, Lei *et al.* applied these criteria to two sweeps (repeats) of the staircase protocol whilst we perform four staircase sweeps in total: two before any other protocols, and two after all other protocols (as shown in Figure 4). Because of these extra repeats of the *staircase* protocol, we apply each QC criterion twice, to both the first two sweeps, and the final two sweeps of the *staircase* protocol.

The first of these criteria that is applied to the time-series data is **QC2**, which ensures that the data has a high signal-to-noise ratio. The noise, $\sigma$, is estimated from the first 200 observations (where the current is assumed to be stationary) by taking the standard deviation of these observations. We then compute the standard deviation of the entire trace and ensure that this is five times larger than our estimated noise. This condition is checked for both the raw trace, and the drug-subtracted, leak-corrected trace. If these criteria are not satisfied, this may be due to an unexpectedly small current, or excessive noise in the trace. A small current may be due to insufficient expression of hERG1a in the cell. Whereas excessive noise may indicate a failure of the experimental equipment (faulty electrodes, for example).

Then, **QC3** ensures that subsequent sweeps result in similarly currents. A large difference between the size of raw traces may indicate that something has changed through time whilst the experiment progressed—a complete cell detachment, for example. Following Lei *et al.*, we retain only data where,

$$\frac{1}{\sqrt{n_{\text{obs}}}}\|\mathbf{y}_1 - \mathbf{y}_2\| < 0.2 \times \frac{1}{2n_{\text{obs}}}(\text{std}(\mathbf{y}_1) + \text{std}(\mathbf{y}_2)), \tag{B.5}$$

for every trace, where $\mathbf{y}_1$ and $\mathbf{y}_2$ are the two time-series traces that are being compared. This check is performed for raw traces (**QC3.raw**), post-drug traces (**QC3.drug**) and leak-corrected, drug-subtracted traces (**QC3.subtracted**).

Next, for **QC5**, we ensure that the drug subtraction has resulted in a significant reduction in current. This is done by first checking that the maximal current (observed during the second half of the protocol) decreases after the addition of dofetilide (**QC5.staircase**), and then checking whether $2\|I_{\text{after}}\|_2\| < I_{\text{before}}\|_2$ (**QC5.1.staircase**).

For **QC6**, we ensure that a positive leak-corrected, drug-subtracted current is observed during +40mV steps. This is done by first checking the three +40mV steps for each sweep of the *staircase* protocol (**QC6.subtracted**, **QC6.1.subtracted** and **QC6.2.subtracted**).

In addition to Lei et al.'s criteria, we check the size of residual, non-leak currents, comparing their size to the pre-drug trace by computing,

$$R_{\text{leftover}} = \frac{\text{RMSE}(I_{\text{obs}}^{(\text{after})}, I_{\text{L}}^{(\text{after})})}{\text{RMSE}(I_{\text{obs}}^{(\text{before})}, I_{\text{L}}^{(\text{before})})}. \tag{B.6}$$

For **QC.R_leftover**, small values of $R_{\text{leftover}}$ indicate that the post-drug trace consists entirely of leak. We then check that,

$$R_{\text{leftover}} < 0.25, \tag{B.7}$$

and denote this criterion **QC.R_leftover**.

**Application of quality-control criteria to other protocols** After applying Lei *et al.*'s QC to their *staircaseramp* protocol, we can apply a subset of these criteria to the other protocols. In particular, protocols **QC1** and **QC4** do not rely on our time-series data, and so, can be applied to each sweep of data; **QC6.subtracted** concerns only the first +40mV step in the protocol, until which point all protocols are identical. We denote the application of these criteria to the remaining protocols by **QC1.all**, **QC4.all** and **QC6.all**, respectively.

**Reversal potential quality-control criteria** We also include some additional checks concerning the apparent reversal potential of the current, $E_{\text{obs}}$ (which we infer as described previously). Having done so, we ensure that $E_{\text{obs}}$ lies in the range covered by our reversal ramp,

$$-110\text{mV} < E_{\text{obs}} < -70\text{mV}, \tag{B.8}$$

removing any wells with a sweep resulting in a $E_{\text{obs}}$ lying outside this range.

Next, we consider the values of $E_{\text{obs}}$ taken from each sweep from a given well, ensuring that the spread in these observed reversal is no greater than 10mV. In other words, we ensure that,

$$\max_i \left\{E_{\text{obs}}^{(i)}\right\} - \min_i \left\{E_{\text{obs}}^{(i)}\right\} < 5\text{mV}, \tag{B.9}$$

where $E_{\text{obs}}^{(i)}$ denotes the reversal potential estimated from a particular sweep, $i$, from the well under consideration. We denote this criterion **QC.Erev.spread**.

**Ensuring stability throughout the experiment** Finally, we apply **QC3.subtracted** to compare the very first and very last repeats of the *staircase* protocol. This, again, is done to ensure that the experimental conditions have not changed significantly over the course of the experiment.

As in [26], all QC criteria are evaluated after the removal of 'capacitive spikes' from the data. In particular, we zero the data observed no more than 2ms after each discontinuity in $V_{\text{cmd}}$. This is because there are large current spikes after each voltage 'step' resulting from the capacitive effect of the cell membrane, which violates our *ideal patch-clamp* assumptions during a short period of time after each voltage step.

**Success rate** After applying these criteria, we discard all but 8 of our 384 wells. This represents a $\approx 2\%$ success rate. This success rate may be improved by instead using more lenient criteria. However, our criteria have been chosen as to leave only high-quality time-series data, with minimal data pollution times-series data, which exhibits consistency over the duration of the experiment. Nevertheless, our approach ensures that our data is suitable for the direct application of our ODE-based models, and the cross-validation methodology introduced in the main text.

| Label | protocols | criterion |
|---|---|---|
| **QC1.Rseal**[1] | all | Check $R_{\text{Seal}} \in [0.1, 1000]$ G$\Omega$ |
| **QC1.Cm** [1] | all | Check $C_{\text{m}} \in [1, 100]$ pF |
| **QC1.Rseries** [1] | all | Check $R_{\text{Series}} \in [1, 25]$ M$\Omega$ |
| **QC2.raw** [1] | *staircase* | Check $\frac{\text{std}(I_{\text{raw}})}{\sigma} > 5$ |
| **QC2.subtracted** [1] | *staircase* | Check $\frac{\text{std}(I_{\text{subtracted}})}{\sigma} > 5$ |
| **QC3.raw** [1] | *staircase* | Check Condition (B.5) with $I^{(1)}_{\text{obs}}, I^{(2)}_{\text{obs}}$ |
| **QC3.drug** [1] | *staircase* | Check Condition $I^{(1)}_{\text{drug}}, I^{(2)}_{\text{drug}}$ |
| **QC3.subtracted** | *staircase* | Check Condition (B.5) $I^{(1)}_{\text{subtracted}}, I^{(2)}_{\text{subtracted}}$ |
| **QC3.bookend** [1] | *staircase* | Check Condition (B.5) with $I^{(1)}_{\text{subtracted}}, I^{(4)}_{\text{subtracted}}$ |
| **QC4.Cm** [1] | all | Check $C^{(\text{before})}_{\text{m}}$ and $C^{(\text{after})}_{\text{m}}$ satisfy Condition (B.4) |
| **QC4.Rseal** [1] | all | Check $R^{(\text{before})}_{\text{seal}}$ and $R^{(\text{after})}_{\text{seal}}$ satisfy Condition (B.4) |
| **QC4.Rseries** [1] | all | Check $R^{(\text{before})}_{\text{series}}$ and $R^{(\text{after})}_{\text{series}}$ satisfy Condition (B.4) |
| **QC5.staircase** [1] | staircase | Check $0.25 \times \max I_{\text{raw}} > \max I_{\text{drug}}$ |
| **QC5.1.staircase** [1] | staircase | Check that $2\|I_{\text{after}}\|_2\| < \|I_{\text{before}}\|_2$ |
| **QC6.subtracted** [1] | all | $\max\{I^{(+40\text{mV})}_{\text{subtracted}}\} > -2\sigma$ where $I^{(+40\text{mV})}_{\text{subtracted}}$ is the current during the first +40mV |
| **QC6.1.subtracted** [1] | *staircase* | $\max\{I^{(+40\text{mV})}_{\text{subtracted}}\} > -2\sigma$ where $I^{(+40\text{mV})}_{\text{subtracted}}$ is the current during the second +40mV step |
| **QC6.2.subtracted** [1] | *staircase* | $\max(I^{(+40\text{mV})}_{\text{subtracted}}) > -2\sigma$ where $I^{(+40\text{mV})}_{\text{subtracted}}$ is the current during the third +40mV step |
| **QC.Erev** | all | Check that $-120\text{mV} < E_{\text{obs}} < -50\text{mV}$ |
| **QC.Erev.spread** | all | Check that $E_{\text{obs}}$ varies by less than 5mV across all repeats and all protocols |
| **QC.R_leftover** | *staircase* | Check that $R_{\text{leftover}} < 0.25$ (Equation (A.4)) |

Table B.1: Quality-control criteria used to process our data. The central column shows which protocols are applied only to the *staircase* protocol, and which criteria are applied to all protocols. Note that **QC3.subtracted** is applied to the first and second sweeps of the *staircase* protocol, as well the third and fourth sweeps. Whereas, **QC3.bookend** is identical but applied to the first and fourth sweeps. However, **QC3.raw**, **QC3.drug**, **QC6.1.subtracted** and **QC6.2.subtracted** are applied only to the first two sweeps and the last two sweeps of the *staircase* protocol.

### C Parameter space boundaries and distributions

To constrain our kinetic rate parameters, we use the same kinetic-rate constrains used in [22]. In particular, for rates of the form $k(V) = a\exp\{bV\}$, we require that,

$$10^{-7}\text{ms}^{-1} < a < 10^{5}\text{ms}^{-1}\,, \tag{C.1}$$

$$10^{-7}\text{mV}^{-1} < b < 10^{5}\text{mV}^{-1}\,, \tag{C.2}$$

$$k(V) < 10^{3}\text{ms}^{-1} \text{ for } \textit{all } V \in [-120\text{mV}, +60\text{mV}] \tag{C.3}$$

and

$$k(V) > 1.67 \times 10^{-5}\text{ms}^{-1} \text{ for } \textit{some } V \in [-120\text{mV}, +60\text{mV}]. \tag{C.4}$$

When calibrating our models with real data, we expect the maximal conductance, $g_{\mathrm{Kr}}$ to vary between cells. This is because we expect the maximal conductance to be roughly proportional to the number of channels, which varies between cells. Likewise, we may expect our maximal conductance to vary if we were to use different cells for our experiments. For this reason, we use maximal conductance constraints that are based on the size of the observed current.

In particular, we consider the minimum observation from the first $-120\,\mathrm{mV}$ step of our leak-corrected and drug-subtracted trace (discarding the first $5\,\mathrm{ms}$), and denote this by $I_{\mathrm{max}}$. We then see that, according to our model,

$$I_{\mathrm{max}} = \bar{g} x_{\mathrm{O}} (V - E_{\mathrm{Kr}}), \tag{C.5}$$

for some unknown proportion of open states, $x_{\mathrm{O}}$ and so, by computing,

$$\bar{g} x_{\mathrm{O}} = \frac{I_{\mathrm{max}}}{V - E_{\mathrm{Kr}}}, \tag{C.6}$$

we can obtain a rough estimate of the maximal conductance, $\bar{g}$. This is possible under the assumption that $0.01 \leq x_{\mathrm{O}} \leq 1$, from which we obtain the constraint,

$$\bar{g}_{\mathrm{min}} \ \leq \ \bar{g} \leq 100 I_{\mathrm{max}} = \bar{g}_{\mathrm{max}}, \tag{C.7}$$

where $\dfrac{I_{\mathrm{max}}}{(V - E_{\mathrm{Kr}})}$ and $\bar{g}_{\mathrm{max}} = 100 I_{\mathrm{max}}$.

**Randomised initial guesses** Following [22] we perform each optimisation multiple times using different initial guesses. Here, we use 30 initial guesses for each sweep of each protocol. Our initial guesses for our '$A$' and '$b$' parameters are sampled from our parameter space using log-uniform and uniform distributions, respectively,

$$\log_{10}\{A_i\} \sim U(-7, 5), \tag{C.8}$$

$$b_i \sim U(10^{-7}, 10^5), \tag{C.9}$$

where $A_i$ and $b_i$ are parameters which characterise some transition rate, $k_i = A_i \exp\{\pm b_i V_m\}$. Unlike [22], we also sample an initial guess for our maximal conductance parameter, $\bar{g}$. Our initial guesses for the maximal conductance, $\bar{g}$, are sampled according to a log-uniform distribution over our parameter space,

$$\log\{\bar{g}\} \sim U(\log\{\bar{g}_{\mathrm{min}}\}, \log\{\bar{g}_{\mathrm{max}}\}), \tag{C.10}$$

where $\bar{g}_{\mathrm{min}}$ and $\bar{g}_{\mathrm{max}}$ are defined as above.

### D Modelling the reversal potential

In Section 5, we assumed that any discrepancy between the observed reversal potential, $E_{\mathrm{obs}}$, and the Nernst potential, $E_{\mathrm{Nernst}}$, was due to some systematic voltage offset. As shown in Figure D.1, this discrepancy is significant. For this reason, we fitted our models under the assumption of a systematic voltage offset, $V_{\mathrm{off}}$, for which we assume $V_{\mathrm{off}} = E_{\mathrm{Nernst}} - E_{\mathrm{obs}}$. In this section, we introduce two other approaches. Firstly, we assume no such voltage offset exists, simply ignoring this discrepancy and assuming that $E_{\mathrm{Kr}} = E_{\mathrm{Nernst}}$ (despite evidence to the contrary shown in Figure D.1). We refer to these assumptions as Case I. Then, in Case II, we assume that $E_{\mathrm{Kr}} = E_{\mathrm{obs}}$, but $V_{\mathrm{off}} = 0$. Finally, in Case III (the case discussed in the main text), we assume $E_{\mathrm{Kr}} = E_{\mathrm{obs}}$ and $V_{\mathrm{off}} = E_{\mathrm{Nernst}} - E_{\mathrm{Kr}}$, such that the discrepancy in the reversal potential is perfectly explained by our voltage offset. In practice, Cases II and III are quite similar because in both cases the current reverses when $V_{\mathrm{m}} = E_{\mathrm{obs}}$, and $E_{\mathrm{obs}}$ varies little over the course of the experiment (see Figure D.1). In fact, we obtain our Case III parameters by fitting under the assumption of Case II, and the voltage-dependent parameters are adjusted accordingly. Cases I, II, III are summarised in Table D.1.

The cases summarised in Table D.1 in different model predictions. The accuracy of these predictions is compared in Figure D.2. Here, we can see that the difference in predictive accuracy between

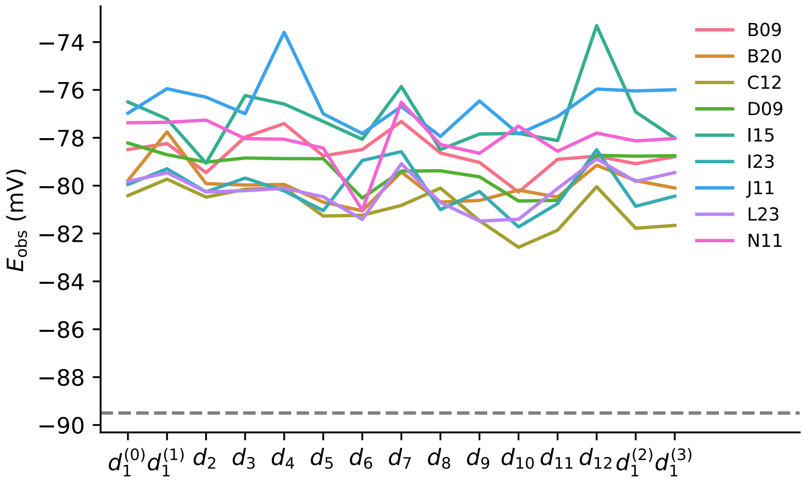


Figure D.1: The observed reversal potential, $E_{obs}$, varies both over the duration of the experiment and between wells. There is a noticeable difference between the observed reversal potentials, $E_{obs}$, and the Nernst potential (shown by the horizontal, grey, dashed line).

| Case | Parameterisation | Physical interpretation |
|---|---|---|
| I | $E_{Kr} = E_{Nernst}$ and $V_m = V_{cmd}$ | The current reverses at the Nernst potential. We fit the model treating $E_{Nernst}$ as a known constant calculated from the Nernst equation. |
| II | $E_{Kr} = E_{obs}$ and $V_m = V_{cmd}$ | $E_{Kr} \neq E_{Nernst}$ (e.g. due to temperature/solution changes). We interpolate $E_{obs}$ from the reversal ramp and treat this as a known constant during model fitting. |
| III | $E_{Kr} = E_{Nernst}$ and $V_m = V_{cmd} + V_{off}$ | $E_{Nernst} \neq E_{obs}$ due to a systematic voltage offset $V_{off}$. We fit the model with the known constant $V_{off} = E_{Nernst} - E_{obs}$ (where $E_{obs}$ is inferred from the reversal ramp as in Case II). |

Table D.1: Different assumptions regarding the models' reversal potential, $E_{Kr}$, and the observed reversal potential, $E_{obs}$, and how this affects model calibration. Apart from the voltage offset in Case III, the patch clamp setup is always treated as *ideal* (no series resistance etc. or time delays in clamp application, and recorded current is solely $I_{Kr}$).

Case I and Case II/III is much more noticeable than any difference between model structures—that is, every model structure produces inaccurate predictions (relatively speaking) when $E_{\mathrm{Kr}} \neq E_{\mathrm{obs}}$. Whilst there is little difference between Cases II and III in this regard, the same is not true of the variability of the resulting parameter estimates. The linear statistical model likelihoods and likelihood ratios presented in the main text are shown for each case in Table D.2. Here, the impact of $V_{\mathrm{off}}$ on the parameter estimates of our voltage-dependent transition rates is noticeable, affecting the apparent well- and protocol-dependence thereof.

| Model | Case | $\mathcal{M}_0$ | $\mathcal{M}_{\mathrm{w}}$ | $\mathcal{M}_{\mathrm{d}}$ | $\mathcal{M}_{\mathrm{w,d}}$ | **LLD-W** | **LLD-D** |
|---|---|---|---|---|---|---|---|
| C-O-I | Case I | 1081.9 | 1172.0 | 1301.0 | 1467.0 | 166.0 | 295.0 |
| | Case II | 1451.5 | 1657.0 | 1709.7 | 2104.7 | 395.1 | 447.7 |
| | Case III | 1465.6 | 1668.5 | 1724.4 | 2111.1 | 386.7 | 442.6 |
| Beattie | Case I | 1231.2 | 1361.5 | 1469.3 | 1723.4 | 254.1 | 362.0 |
| | Case II | 1547.4 | 1804.0 | 1757.0 | 2219.3 | 462.3 | 415.4 |
| | Case III | 1563.3 | 1817.0 | 1784.2 | 2247.2 | 462.9 | 430.2 |
| Kemp | Case I | 118.9 | 1220.1 | 1486.5 | 1671.3 | 184.8 | 451.2 |
| | Case II | 914.7 | 1081.8 | 1096.0 | 1337.2 | 241.2 | 255.4 |
| | Case III | 931.8 | 1099.8 | 1113.7 | 1356.3 | 242.6 | 256.5 |
| Wang | Case I | −1919.8 | −1851.5 | −1578.2 | −1474.3 | 103.9 | 377.1 |
| | Case II | −506.4 | −379.0 | −216.6 | 14.6 | 202.0 | 364.4 |
| | Case III | −2082.1 | −1976.1 | −1836.7 | −1673.5 | 163.2 | 302.6 |

Table D.2: Log-likelihoods and log-likelihood differences (LLD-W and LLD-D) for each of the linear models, applied to all of our models (that is, the collection of parameter estimates obtained using each model). Here we see that in each case, and for each model, that both well- and protocol-effects very significant. Whilst **LLD-D** suggests that there is discrepancy between the recordings of $\mathrm{I}_{\mathrm{Kr}}$ (taken from each well) and the dynamics of our models, the significance of **LLD-W** may suggest the presence of latent, well-dependent effects.

### E Summary of model residuals and predictive discrepancy

**Model fitting residuals** As described in Section 5, we may summarise the ability of our models to recapitulate our recordings of $\mathrm{I}_{\mathrm{Kr}}$ by computing and averaging model residuals. Such an example is shown for the Beattie model in Figure 7 of the main article. In this section we, provide similar figures, Figures E.1, E.2 and Figures E.3, for the C-O-I, Kemp and Wang models, respectively.

**Ensemble prediction scores** Similarly, we may reproduce Figure 8 for the remaining models, allowing a comparison of our predictive models and their tendency to produce discrepant prediction. Figures E.4, E.5 and E.6 show the $T$ statistic (introduced in Section 5) averaged across wells for each time point of each protocol for the C-O-I model, Kemp model [29] and Wang model [28], respectively.

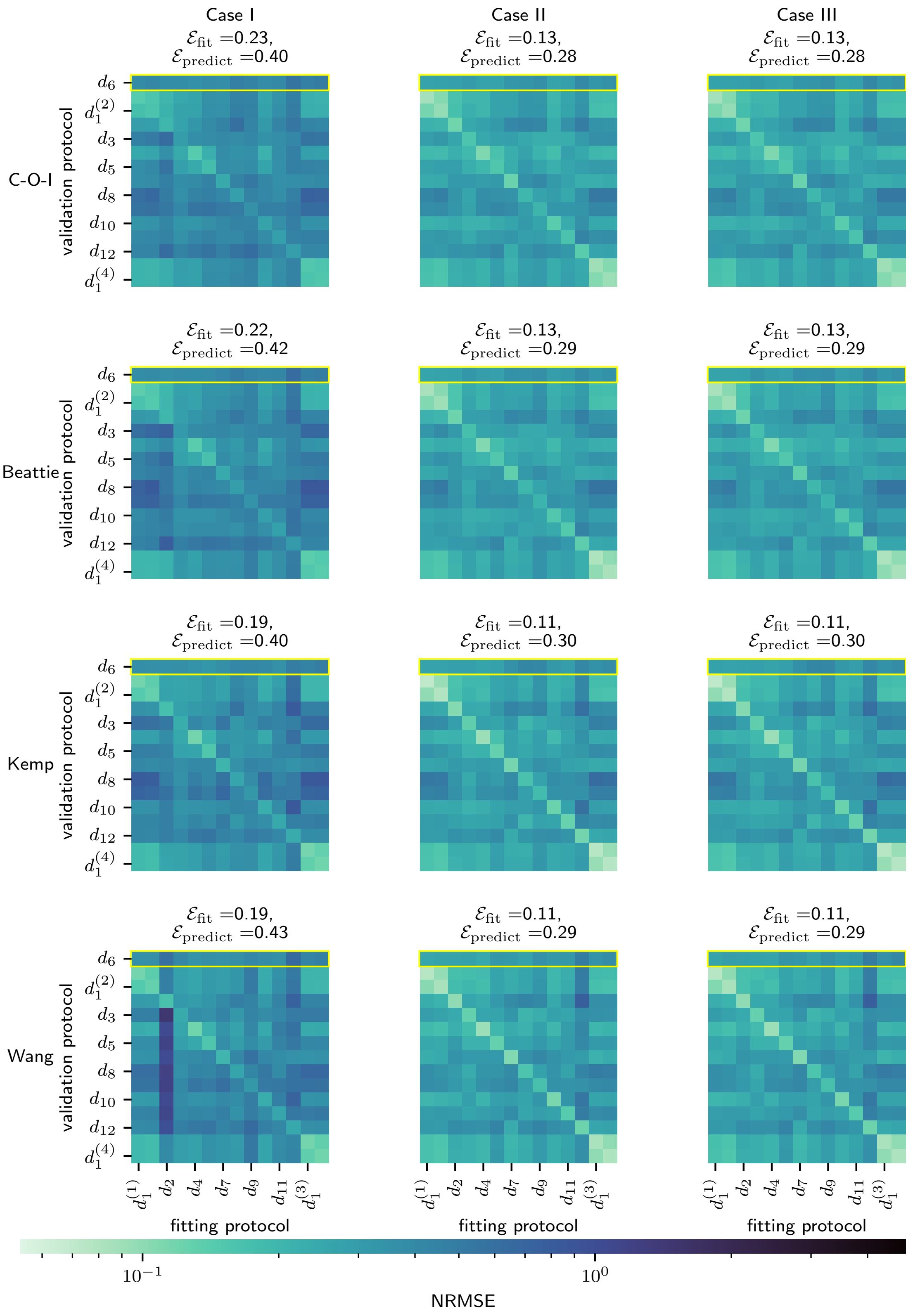


Figure D.2: A comparison of the predictive performance of our chosen model structures (under Cases I, II and III). Each heatmap shows the average normalised root-mean-square error when a particular model is trained and validated using a particular pair of protocols.

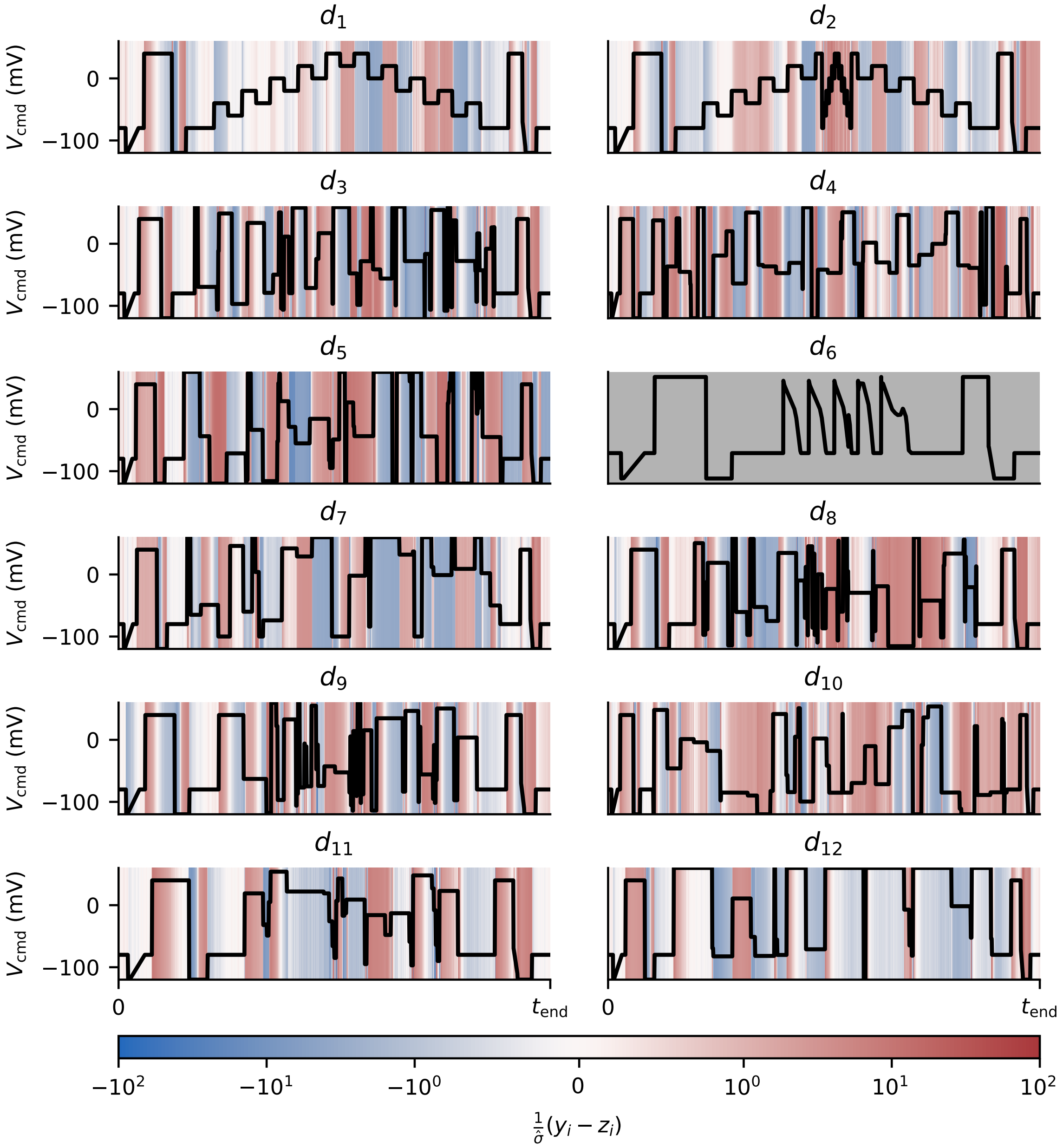


Figure E.1: The C-O-I model is seemingly unable to recapitulate certain sections of our training protocols. Each panel shows the average residual when the C-O-I model is using data from each protocol, down-weighted by our estimate of the standard deviation of the noise. These values are clipped between −100 and +100. Protocol $d_6$, which is used only for validation and not fitting, is shown in grey.

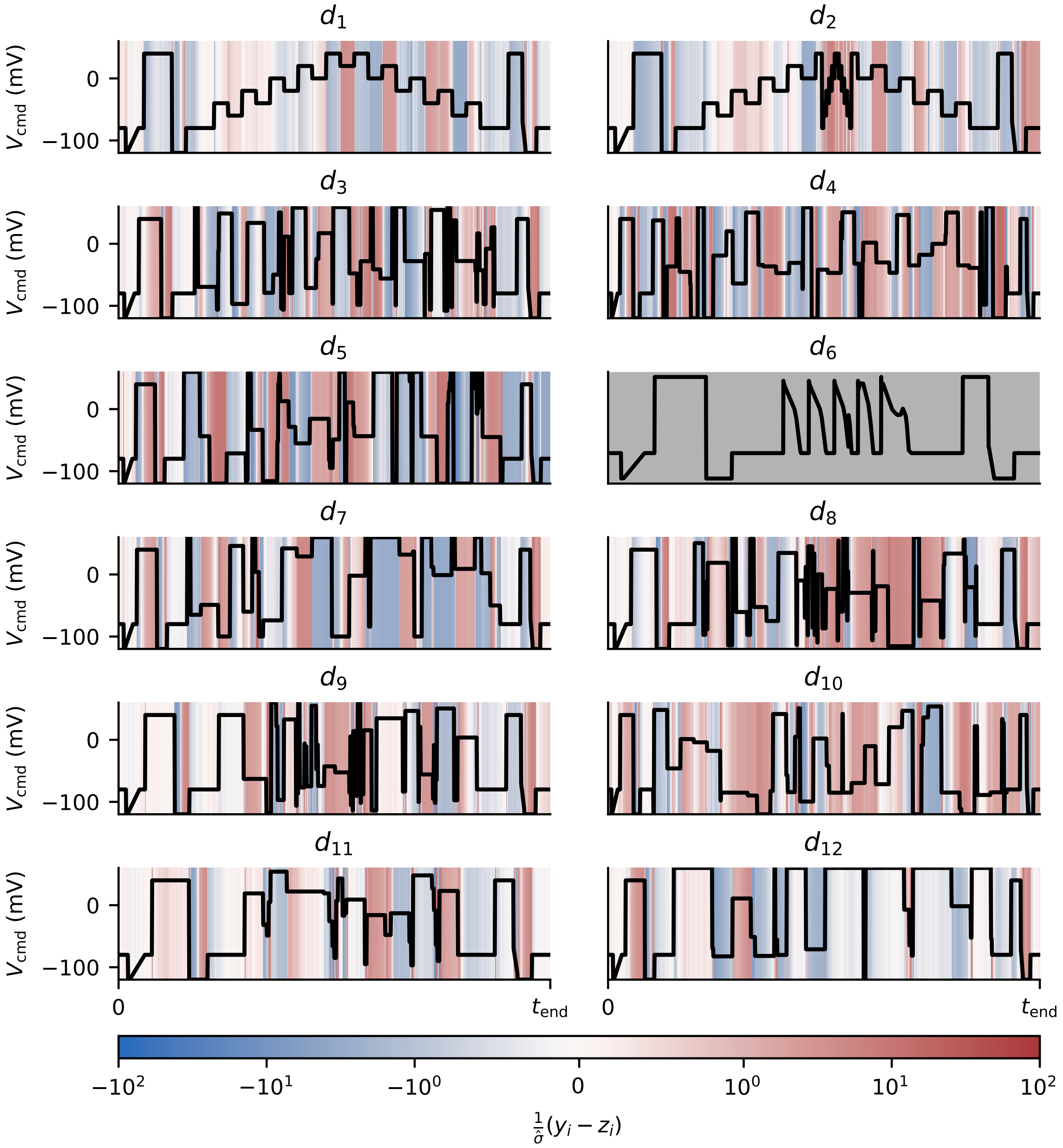


Figure E.2: The Kemp model is seemingly unable to recapitulate certain sections of our training protocols. Each panel shows the average residual when the Wang model is fitted using data from each protocol, down-weighted by our estimate of the standard deviation of the noise. These values are clipped between −100 and +100. Protocol $d_6$, which is used only for validation and not fitting, is shown in grey.

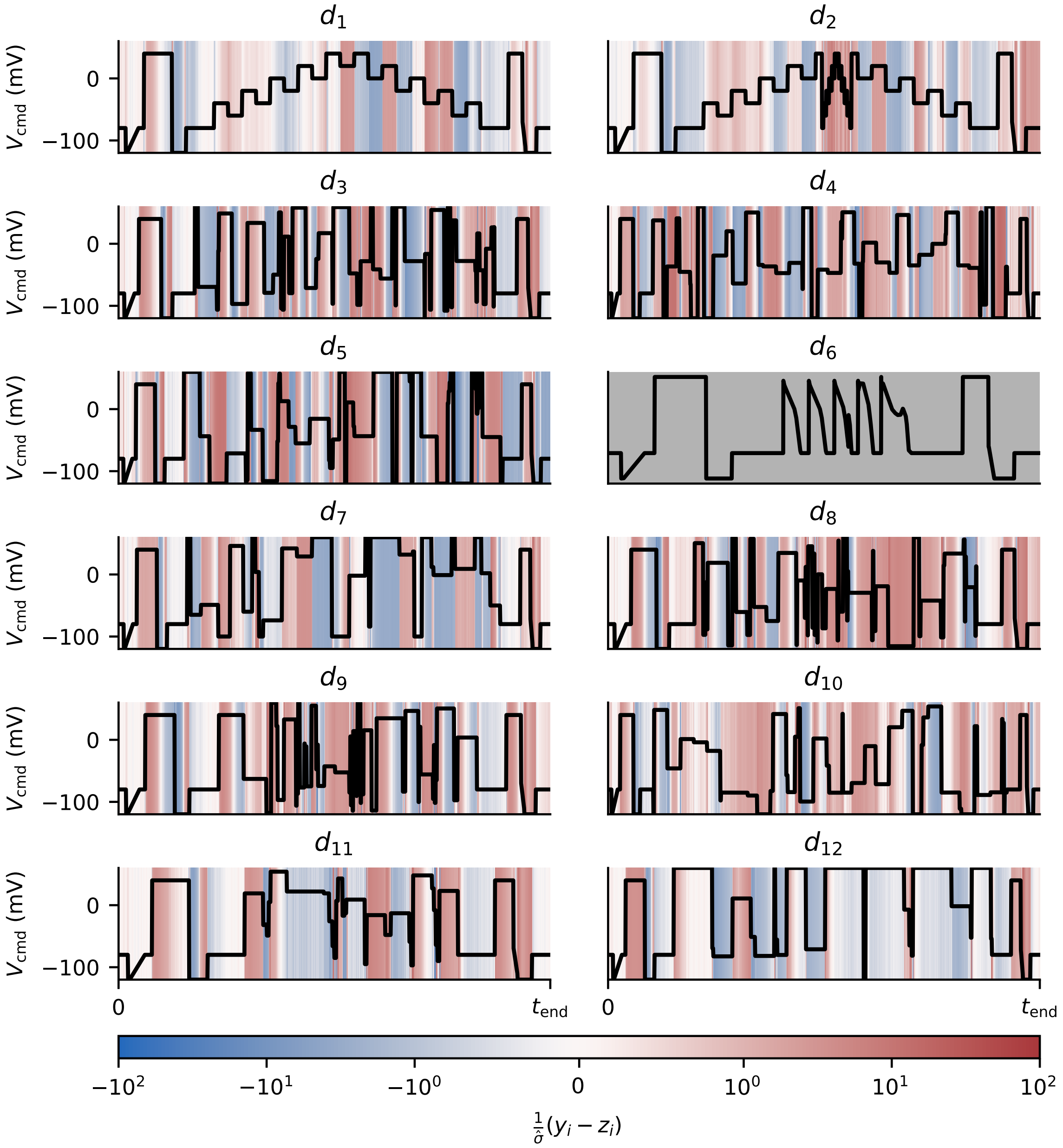


Figure E.3: The Wang model is seemingly unable to recapitulate certain sections of our training protocols. Each panel shows the average residual when the Wang model is using data from each protocol, down-weighted by our estimate of the standard deviation of the noise. These values are clipped between −100 and +100. Protocol $d_6$, which is used only for validation and not fitting, is shown in grey.

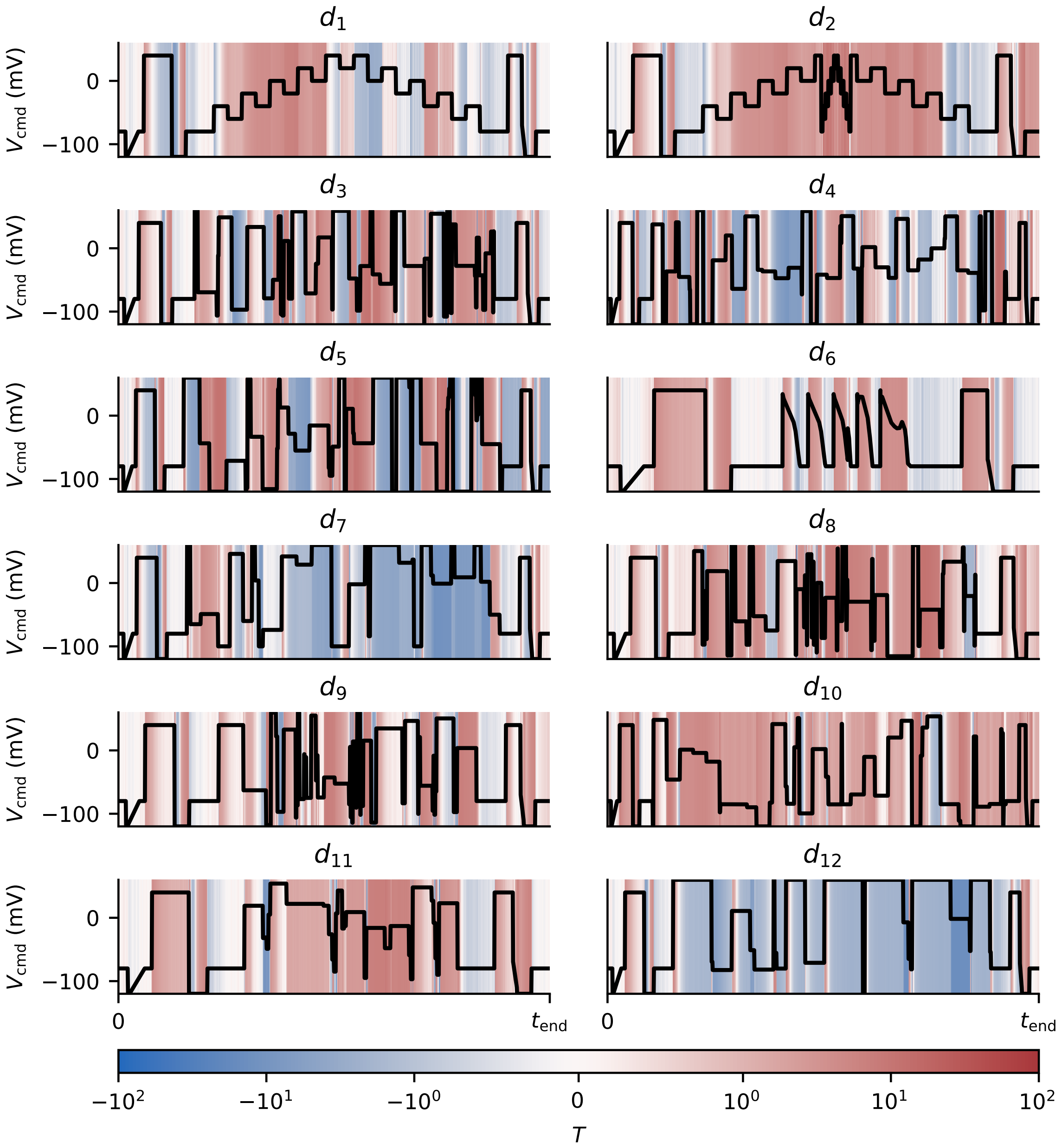


Figure E.4: A summary of the predictions of the C-O-I model. When our ensemble of predictions consistently overestimates $I_{Kr}$, this is shown by positive values of $T$. Similarly, consistent overestimation is shown by negative values of $T$. Values of $T$ are clipped between $-100$ and $+100$.

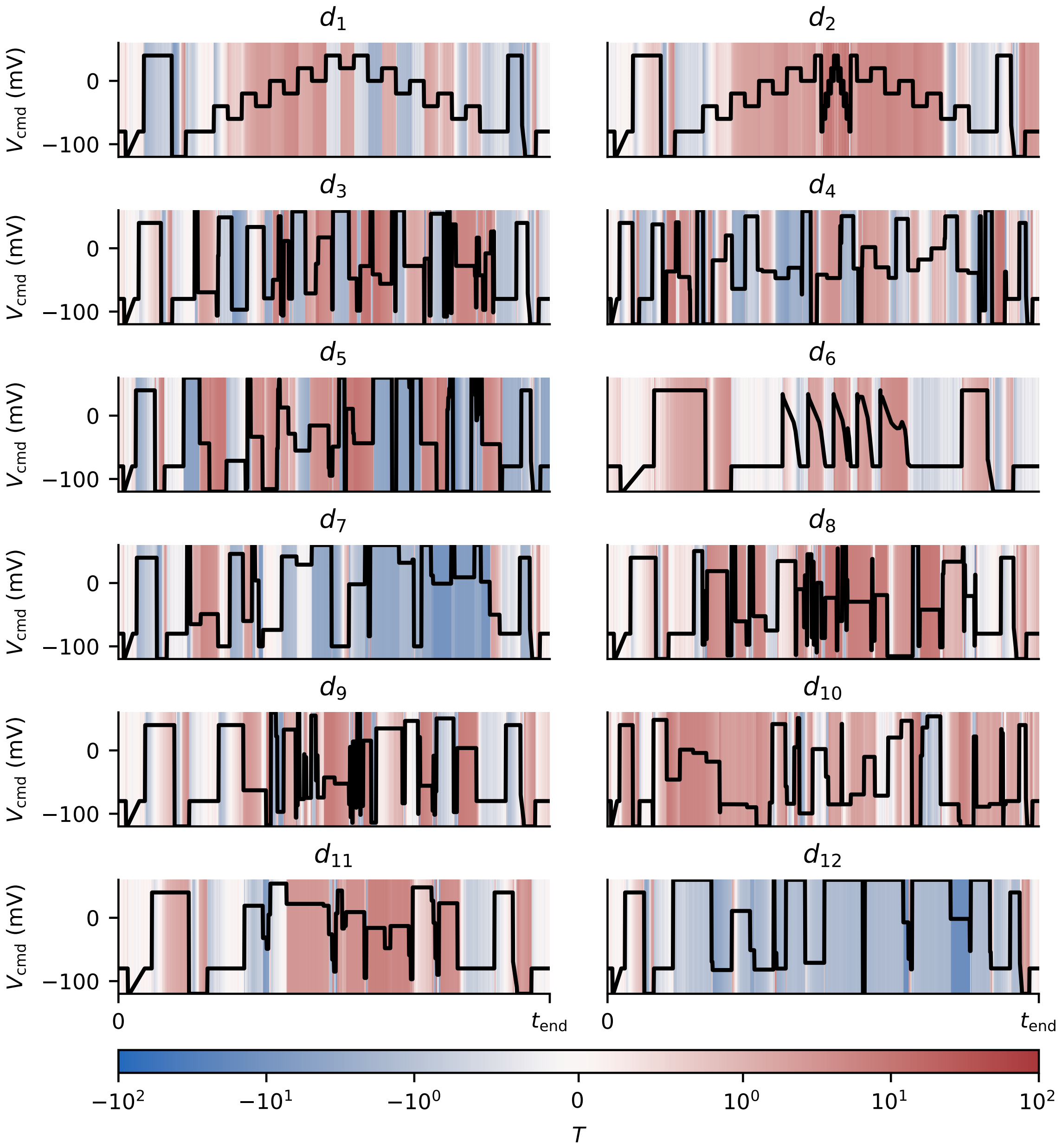


Figure E.5: A summary of the predictions of the Kemp model. When our ensemble of predictions consistently overestimates $I_{Kr}$, this is shown by positive values of $T$. Similarly, consistent overestimation is shown by negative values of $T$. Values of $T$ are clipped between $-100$ and $+100$.

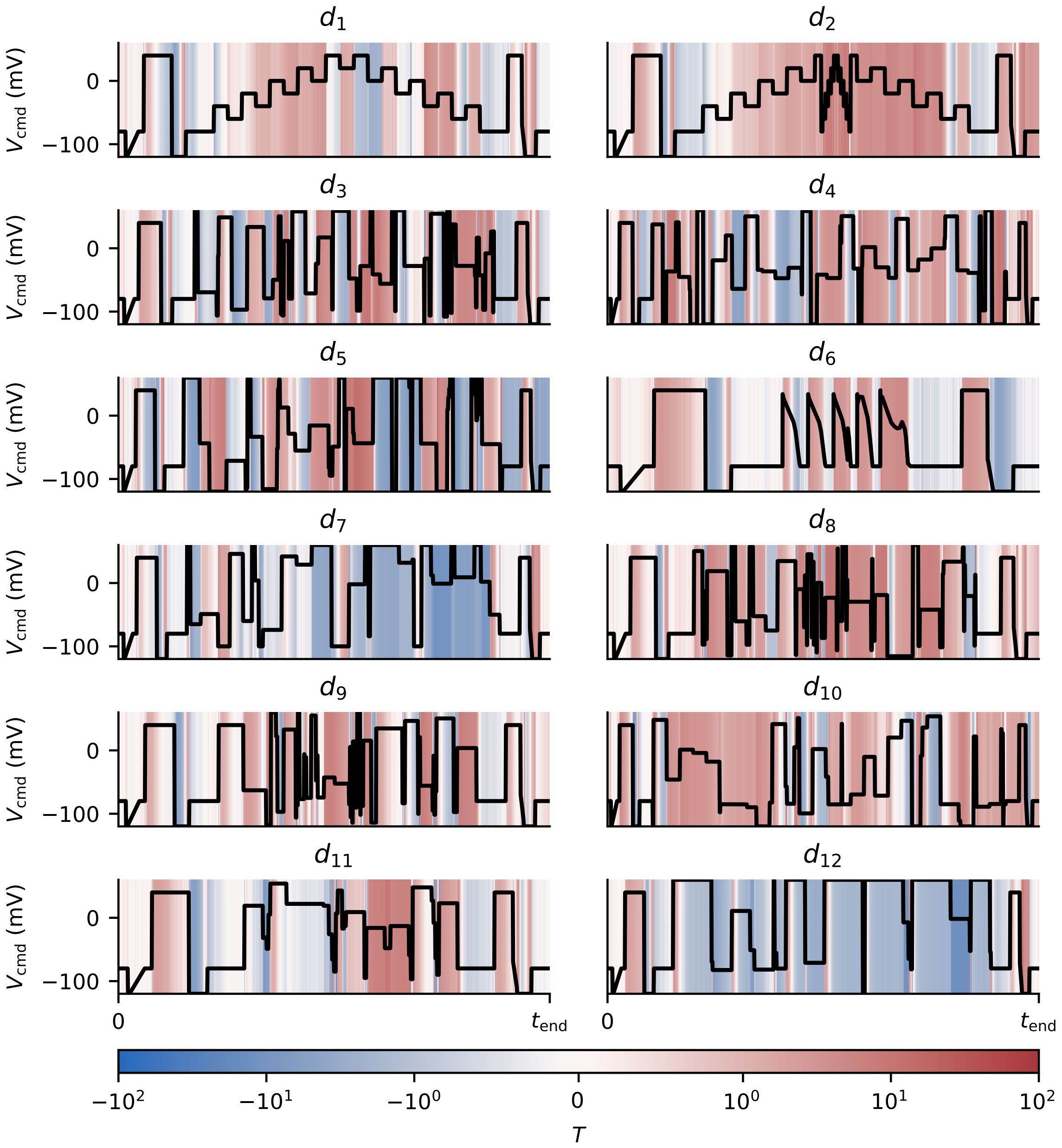


Figure E.6: A summary of the predictions of the Wang model. When our ensemble of predictions consistently overestimates $I_{Kr}$, this is shown by positive values of $T$. Similarly, consistent overestimation is shown by negative values of $T$. Values of $T$ are clipped between $-100$ and $+100$.
